# Relapse-like behavior and nAChR sensitization following intermittent access nicotine self-administration

**DOI:** 10.1101/2022.01.07.475355

**Authors:** Melissa A. Tapia, Xiao-Tao Jin, Brenton R. Tucker, Leanne N. Thomas, Noah B. Walker, Veronica J. Kim, Steven E. Albertson, Naresh Damuka, Ivan Krizan, Seby Edassery, Jeffrey N. Savas, Kiran Kumar Solingapuram Sai, Sara R. Jones, Ryan M. Drenan

## Abstract

Many tobacco smokers consume nicotine intermittently, but the underlying mechanisms and neurobiological changes associated with intermittent nicotine intake are unclear. Understanding intermittent nicotine intake is a high priority, as it could promote therapeutic strategies to attenuate tobacco consumption. We examined nicotine intake behavior and neurobiological changes in male rats that were trained to self-administer nicotine during brief (5 min) trials interspersed with longer (15 min) drug-free periods. Rats readily adapted to intermittent access (IntA) SA following acquisition on a continuous access (ContA) schedule. Probabilistic analysis of IntA nicotine SA suggested reduced nicotine loading behavior compared to ContA, and nicotine pharmacokinetic modeling revealed that rats taking nicotine intermittently may have increased intake to maintain blood levels of nicotine that are comparable to ContA SA. After IntA nicotine SA, rats exhibited an increase in unreinforced responses for nicotine-associated cues (incubation of craving) and specific alterations in the striatal proteome after 7 days without nicotine. IntA nicotine SA also induced nAChR functional upregulation in the interpeduncular nucleus (IPN), and it enhanced nicotine binding in the brain as determined via [^11^C]nicotine positron emission tomography. Reducing the saliency of the cue conditions during the 5 min access periods attenuated nicotine intake, but incubation of craving was preserved. Together, these results indicate that IntA conditions promote nicotine SA and nicotine seeking after a nicotine-free period.

## INTRODUCTION

Tobacco dependence is a leading cause of preventable death, and the economic costs associated with tobacco usage are staggering. For example, tobacco addiction in the United States costs >$300 billion in lost wages/productivity and healthcare costs. Approximately 1,300 Americans die *each day* from tobacco-related disease (DHHS, 2014). Clearly, developing new tobacco cessation strategies and identifying mechanisms giving rise to tobacco consumption is an important scientific and public health priority.

Among several research systems available for modeling human tobacco usage, rodent self-administration (SA) models have contributed important insights into tobacco addiction mechanisms (Corrigall and Coen, 1989, 1991; Corrigall et al., 1994; Donny et al., 1995; Donny et al., 1999; Donny et al., 2000). Moreover, these models have helped validate drugs such as varenicline (Rollema et al., 2007), which is modestly efficacious as a tobacco cessation therapeutic. Rat models of nicotine SA have largely employed fixed-ratio schedules of reinforcement, often with short (1-2 h) sessions, an FR1 to FR5 response requirement, and a timeout (usually 20 s) period after an infusion is earned (Caille et al., 2012). However, these parameters may not fully capture the nuances of human nicotine intake. Daily cigarette smokers smoke more during morning hours than during other times of the day (Shiffman et al., 2014). Smokers structure their activities around designated times for nicotine intake, such as smoke breaks during work hours and social activities in the evening and on weekends (Shiffman et al., 2014). In many smokers, this inevitably leads to nicotine intake patterns that are intermittent; nicotine intake will rise and fall numerous times per day.

Intermittent drug intake patterns, where short periods of drug access are interspersed with longer periods without access within a single session (Morgan et al., 2002), have only recently been appreciated for their ability to promote drug-taking and produce distinct neurobiological changes (Kawa et al., 2019). When rats self-administer cocaine on a schedule that results in rapid spikes in brain levels of cocaine (intermittent access; IntA), motivation to maintain those levels is enhanced compared to a long access (LgA) SA schedule that maintains high levels of cocaine in the brain (Zimmer and Roberts, 2012; Kawa et al., 2016). Consistently, dopamine (DA) release and DA uptake exhibited tolerance following LgA cocaine SA but *sensitization* following IntA cocaine SA (Calipari et al., 2013).

Although there are no identifiable reports in the literature involving nicotine SA sessions with short access periods and intervening periods without access, intermittent nicotine exposure has been examined. A paradigm involving cycles of 4 days on nicotine SA and 3 days off, where the nicotine dose was increased each cycle, led to greater nicotine intake versus continuous access to a single dose (O’Dell and Koob, 2007). When the cycle was 1 day on nicotine SA and 2-3 days off, 23 h access led to more profound anhedonia compared to 1 h access sessions (Geste et al., 2020). Moreover, non-contingent intermittent (12 h on, 12 h off) nicotine exposure induced an increase in nAChR binding sites in several brain areas (Semenova et al., 2018). Although these examples of nicotine exposure have aspects of intermittency and suggest that variation in the timing of nicotine delivery induces unique changes to physiology and/or behavior, we could not identify research using IntA that parallels the recent work conducted with cocaine.

Here, we examined nicotine SA behavior during sessions with 5 min access periods and 15 min intervening periods without access to determine 1) whether rats participate in nicotine SA under these conditions, 2) how their nicotine intake compares to continuous access, 3) whether intermittent access impacts relapse-like behavior, and 4) if this access paradigm is sufficient to produce sensitization of nicotinic acetylcholine receptors (nAChRs) in relevant brain areas.

## MATERIALS AND METHODS

### Materials

Nicotine hydrogen tartrate salt was obtained from Glentham Life Sciences (catalog #GL9693-5G). Injectable heparin sodium (catalog #07-892-8971) and injectable meloxicam (catalog #07-891-7959) were obtained from Patterson Veterinary Supply. Mecamylamine hydrochloride was obtained from Sigma-Aldrich (catalog #M9020).

### Animals

All experimental protocols involving rats were reviewed and approved by the Wake Forest University Institutional Animal Care and Use Committee. Procedures also followed the guidelines for the care and use of animals provided by the National Institutes of Health Office of Laboratory Animal Welfare. All efforts were made to minimize animal distress and suffering during experimental procedures, including during the use of anesthesia. Male Sprague Dawley rats (Envigo; total n = 106) were ∼300 g (approximately eight weeks old) when they arrived at our facility. Rats were housed at 22°C on a reverse 12/12 h light/dark cycle (4 P.M. lights on, 4 A.M. lights off).

### Apparati

Rats were trained in Med Associates operant chambers (interior dimensions, in inches: 11.9 × 9.4 × 11.3, catalog #MED-007-CT-B1) located within sound-attenuating cabinets. The SA system was housed in a dedicated room within the same laboratory suite as the rat’s housing room. A PC computer was used to control the SA system via Med PC IV software. Each chamber had transparent plastic walls, a stainless-steel grid floor, and was equipped on the right-side wall with two nose pokes (2.4 inches from grid floor to nose poke center) which flanked a pellet receptacle coupled to a pellet dispenser. A white stimulus light was located above each nose poke, and a house light was located at the top of the chamber on the left-side wall. During food and drug SA sessions, nose pokes on the active nose poke activated either the pellet dispenser or an infusion pump, respectively.

Nose pokes on the inactive nose poke had no consequence. For intravenous drug infusions, each rat’s catheter was connected to a liquid swivel via polyethylene tubing protected by a metal spring. The liquid swivel was connected to a 10-mL syringe loaded onto the syringe pump.

### Operant Food Training

Approximately one week after arrival, pair-housed rats were food-restricted for several days to enhance their participation in operant responding. Rats were fed standard chow (LabDiet Prolab RMH 3000 5P00, catalog #0001495) (40g per cage) once per day at least 1 h after finishing testing. Water was available *ad libitum* except during operant behavioral sessions. Food training sessions were 1 h in duration, and rats were trained to nose poke for food pellets (45 mg; Bio-Serv Dustless Precision Pellets, catalog #F0021) on the same nose poke that would subsequently be paired with drug infusions in nicotine IVSA sessions. A fixed ratio 1 (FR1; no timeout) schedule was used for food training; no visual cues (stimulus light, house light) were illuminated during the session and rats could earn a maximum of 75 food pellets during the 1-h session. Once each rat successfully earned at least 50 pellets with at least a 2:1 preference for the active nose poke over the inactive nose poke, no further food training was conducted. Rats met this criterion on average between three and five days.

### Indwelling Jugular Catheter Surgery

After acquiring food operant responding, rats were anesthetized with isoflurane (3% induction, 2–3% maintenance) and implanted with indwelling jugular catheters (Instech, catalog #C30PU-RJV1402). Meloxicam (2 mg/kg) was administered postoperatively to relieve pain and reduce inflammation. Rats were singly housed following surgery and throughout all SA procedures. Rats were allowed 7 d for recovery from surgery, and catheters were flushed several times during this recovery period with heparin sodium dissolved in sterile saline.

### Intravenous Drug Self-Administration

After recovery from catheter surgery, rats were allowed to self-administer saline or nicotine (0.03 mg/kg free base/infusion) in a volume of 0.035 mL over 2 s during 2 h SA sessions, Monday through Friday (no SA sessions occurred on weekends). (−)-Nicotine hydrogen tartrate salt (Glentham Life Sciences) was dissolved in sterile saline, and the pH was adjusted to 7.4. Infusions, delivered by an infusion pump, were triggered by one nose poke response on the active nose poke. Infusions (2 s duration) were simultaneously paired with illumination of the stimulus light over the active nose poke for 3 s. An active nose poke response that resulted in an infusion extinguished the house light for a 20-s timeout period (TO-20), during which responding was recorded but had no consequence. Responses on the inactive nose poke were recorded but had no scheduled consequence. At the end of the session, the house light was extinguished and responding had no consequences. Rats were removed from the training chambers as soon as possible after the end of the 2-h session and were rationed to 20 g of standard lab chow at least 1 h after finishing their sessions. Modified chow availability was used throughout self-administration and a range of 85-90% of free-feeding body weight was maintained. Rats were allowed to self-administer saline or nicotine for 10 sessions on a FR1/TO-20 schedule of reinforcement (acquisition). If any one of the following occurred, the rat did not move on to the Intermittent Access (IntA) portion of the study: (1) less than 10 nicotine infusions were earned within the 2-h session for 2 or more consecutive sessions, (2) the ratio of active to inactive nose pokes was less than 2.0 for 3 or more consecutive sessions and (3) a drop of 75% or greater in responding on the active nose poke occurred over the final 5 sessions of acquisition. A total of 10 rats failed to meet the above criteria, with an additional 11 rats excluded due to loss of catheter patency, thus a total of 85 rats completed acquisition.

IntA procedures were similar to those previously described (Zimmer and Roberts, 2012; Kawa et al., 2016). Each session consisted of 12 alternating Drug-Available and No Drug-Available periods lasting 5- and 15- min, respectively. The house light was on at the start of the session. The Drug Available period started immediately after the rats were placed into the chamber. During the Drug Available period, an active nose poke response (FR1/TO-20) resulted in an infusion, for which the house light was extinguished for a 20- s timeout period. During this time, responding was recorded but had no consequence. Responses on the inactive nose poke were recorded but had no scheduled consequence. After the 5-min period, the house light turned off and signaled a 15-min No Drug Available period. During the No Drug Available period, all nose pokes were recorded but had no consequences. The 5- min Drug Available and 15- min No Drug Available periods repeated themselves for a total of 12 Drug Available and 12 No Drug Available periods, resulting in a 4 h session, with 1 h of nicotine availability. Rats self-administered saline or nicotine on the IntA paradigm for 7 sessions.

### Brain Slice Preparation and Recording Solutions

Rats were anesthetized with isoflurane before trans-cardiac perfusion with oxygenated (95% O2/5% CO2), 4°C N-methyl-D-glucamine (NMDG)-based recovery solution that contains (in mM): 93 NMDG, 2.5 KCl, 1.2 NaH2PO4, 30 NaHCO3, 20 HEPES, 25 glucose, 5 sodium ascorbate, 2 thiourea, 3 sodium pyruvate, 10 MgSO4·7H2O, and 0.5 CaCl2·2H2O; 300-310 mOsm; pH 7.3-7.4. Brains were immediately dissected after the perfusion and held in oxygenated, 4°C recovery solution for one minute before cutting a brain block containing the IPN and sectioning the brain with a vibratome (VT1200S; Leica). Coronal slices (250 µm) were sectioned through the IPN and transferred to oxygenated, 33°C recovery solution for 12 min. Slices were then kept in holding solution containing in mM: 92 NaCl, 2.5 KCl, 1.2 NaH2PO4, 30 NaHCO3, 20 HEPES, 25 glucose, 5 sodium ascorbate, 2 thiourea, 3 sodium pyruvate, 2 MgSO4·7H2O, and 2 CaCl2·2H2O; 300-310 mOsm; pH 7.3-7.4 for 60 min or more before recordings. Brain slices were transferred to a recording chamber (1 mL volume), being continuously superfused at a rate of 1.5-2.0 mL/min with oxygenated 32°C recording solution. For our recording chamber and solution flow rate, we estimate that complete solution exchange occurs in 5 to 8 min. The recording solution contained (in mM): 124 NaCl, 2.5 KCl, 1.2 NaH2PO4, 24 NaHCO3, 12.5 glucose, 2 MgSO4·7H2O, 2 CaCl2·2H2O, 0.01 CNQX, 0.03 D-AP5, and 0.1 picrotoxin; 300-310 mOsm; pH 7.3-7.4).

For puffer experiments, the recording solution was supplemented with 1 µM atropine. Patch pipettes were pulled from borosilicate glass capillary tubes (1B150F-4; World Precision Instruments) using a programmable microelectrode puller (P-97; Sutter Instrument). Tip resistance ranged from 7.0 to 10.0 MΩ when filled with internal solution. A potassium gluconate-based internal solution was used for recordings (in mM): 135 potassium gluconate, 5 EGTA, 0.5 CaCl2, 2 MgCl2, 10 HEPES, 2 MgATP, and 0.1 GTP; pH adjusted to 7.25 with Tris base; osmolarity adjusted to 290 mOsm with sucrose. The internal solution contained QX314 (2 mM) for improved voltage control.

### Patch Clamp Electrophysiology

Electrophysiology experiments were conducted using a Nikon Eclipse FN-1 upright microscope equipped with a 40x (0.8 NA) water-dipping (3.3 mm working distance) objective. Neurons in the rostral IPN (IPR) were targeted for recording. Neurons within brain slices were first visualized with infrared or visible differential interference contrast (DIC) optics. A computer running pCLAMP 10 software was used to acquire whole-cell recordings along with a Multiclamp 700B amplifier and a Digidata 1550A A/D converter (all from Molecular Devices Inc.). Data were sampled at 10 kHz and low pass filtered at 1 kHz. Immediately prior to giga seal formation, the junction potential between the patch pipette and the superfusion medium was nulled. Series resistance was uncompensated. To record physiological events following local application of drugs, a drug-filled pipette was moved to within 20-40 µm of the recorded neuron using a second micromanipulator. A Picospritzer (General Valve) dispensed drug (dissolved in recording solution) onto the recorded neuron via a pressure ejection. Pipette location relative to the recorded cell, along with ejection pressure, were held constant throughout the recording. Ejection duration was varied at half-log steps to enable collection of quasi- concentration response curves.

### Radiochemistry

[^11^C]nicotine was produced following published procedures (Garg et al., 2017; Solingapuram Sai et al., 2020). Briefly, [^11^C]MeI from GE-FXC module was bubbled to the reaction vial containing precursor nor-nicotine (∼0.5 mg) in anhydrous DMF (0.15 mL) and ACN (0.35 mL) for ∼5 min at room temperature. After the complete transfer of radioactivity, the sealed reaction vial was then heated at 120°C for 3 min. The reaction mixture was quenched with HPLC mobile phase (0.6 mL) and injected onto a normal reverse phase semi-preparative C18 LUNA (250 × 10 mm, 10 µm) HPLC column to purify [^11^C]nicotine. The isocratic HPLC mobile phase solution consisted of 65% acetonitrile, 35% 0.1 M aqueous ammonium formate buffer solution (pH 6.0–6.5) with UV λ @ 254 nm and a flow rate of 5.0 mL/min. The product [^11^C]nicotine (Rt = 9.0-11.0 min) was collected and diluted with 50 mL of deionized water, and passed through a solid phase extraction cartridge (Waters Sep-Pak C18; Milford, MA) to trap the radioactive product. [^11^C]nicotine was then directly eluted from the cartridge with 10% ethanol in saline into a sterile vial through a sterile 0.2 µm filter for all animal studies.

### microPET Imaging

Rodent microPET/CT imaging of [^11^C]nicotine was performed as described previously (Solingapuram Sai et al., 2018). In brief, anesthetized rats (n = 4 nicotine SA and n = 6 saline SA) were injected with [^11^C]nicotine (200 ± 10 µCi) into the tail vein and a dynamic 0-30 min brain scan was obtained using a TriFoil PET/CT scanner. The DICOM images were reconstructed using attenuation correction and analyzed using πPMOD analysis software.

### Biodistribution Studies

Standard post-PET biodistribution studies (Solingapuram Sai et al., 2012) were performed in the same rats used for microPET. After PET acquisition, rats were euthanized and uptake of radiotracer in the brain was measured using a γ- counter. The uptake was calculated as the percentage of injected dose per gram of tissue (%ID/g).

### Pharmacokinetic Modeling

Nicotine pharmacokinetic simulations in rats were conducted using MATLAB and SimBiology (MathWorks). A two-compartment model was built, simulating a central (blood, plasma, cerebrospinal fluid, etc.) and peripheral (fat, muscle, and other poorly perfused tissues) compartment. Hereafter, “plasma” denotes the central compartment. Model parameters, included in the simulation code (https://github.com/ryandrenan/Tapia_et_al), were selected based on previous studies (Kyerematen et al., 1988; Shivange et al., 2019). Key model parameters include: central compartment capacity (5 L/kg), elimination rate constant (0.8), forward rate constant (central->peripheral; 1.5), reverse rate constant (peripheral->central; 1.2), and clearance rate constant (1.4). Limitations include, but are not limited to, the following: 1) a standard 350 g rat was modeled, and 2) variability in nicotine metabolism across animals was not accounted for. Model parameters were first validated by comparing rat plasma nicotine kinetics following an arterial bolus dose (Kyerematen et al., 1988) to the nicotine kinetics predicted by the model after simulating the same bolus dose into the central compartment. Moreover, the peak plasma [nicotine] levels predicted by the model agree with other reports that measured rat nicotine levels in plasma after intravenous dosing (Craig et al., 2014). To model plasma nicotine levels in our experimental rats, the model was enabled to accept infusion time vectors from SA sessions. The model returned an estimated [nicotine] vs. time profile, which was used to calculate key metrics such as peak plasma [nicotine] and plasma area under the curve (AUC) for specified time periods after the start of the SA session. Estimated steady-state plasma [nicotine] levels (10-50 ng/mL) in our experimental rats during nicotine SA sessions were similar to plasma nicotine levels measured in human smokers during unrestricted cigarette smoking (Benowitz et al., 1983) and during regular, intermittent cigarette smoking (Isaac and Rand, 1972; Benowitz et al., 1982). In particular, we observed peaks and troughs in the plasma nicotine vs. time profile that were reminiscent of data from humans during intermittent cigarette smoking (Isaac and Rand, 1972).

### Proteomic analysis

Rats (n=16) were euthanized and the dorsal striatum was acutely dissected, pooled (i.e. 2 extracts from the same brain), and flash frozen. Proteins were extracted from brain tissue using RIPA buffer with protease inhibitors, followed by purification via chloroform methanol precipitation. The precipitated proteins were solubilized in 6M Guanidine Hydrochloride, reduced with DTT, alkylated using IAA, and digested with lys-C for 3 hours and trypsin overnight at 37°C . The trypsin digest was acidified, and peptides were purified using C18 HyperSep columns. Each tube of purified peptides (100 µg) were labeled with 16 plex TMT pro tags as we have previously reported, fractionated using high pH columns to 8 fractions, and each fraction was analyzed with a 4 h gradient via the MS3 multinotch method (Ting et al., 2011; McAlister et al., 2014; Jongkamonwiwat et al., 2020). Protein identification and quantitation was performed with ProLucid, DTASelect, and Census within the IP2 bioinformatic analysis pipeline (Bruker) (Xu et al., 2015). We used the Uniprot Rat UP000002494 database (downloaded on 3-21-2020). In the search, we considered half- and fully tryptic peptides within 10 ppm of the expected precursor m/z and required only one peptide to be identified/quantified for each protein. The protein false discovery rate was controlled to 1% at the protein level based on the target-decoy strategy. We used the NCBI BRB-ArrayTools for class comparison and mapped each protein group to the corresponding gene (n=4399). We used a univariate test: A two-sample T-test (with random variance model) and report proteins with a false discovery rate (FDR) <0.05. The FDR is an estimate of the proportion of the proteins/genes with univariate *p* values less than or equal to the one in that row that represents false positives (i.e. Benjamini and Hochberg method). The heat map (**Fig. 5D**) was made using dynamic heat map maker within the BRB array tool. GO analysis (including GO dot plot) was executed using clusterProfiler (version 3.99.1) and Enrichplot (version 1.10.2) R packages.

### Experimental Design and Statistical Analysis

SA data files, produced by Med Associates MedPC IV software, were processed and analyzed with custom scripts written in MATLAB and/or GraphPad Prism 9. Electrophysiology data files produced by pClamp were processed with custom MATLAB scripts and Origin (OriginLab) software. Scalable vector graphics were produced from MATLAB figures using functions written by Salva Ardid (https://github.com/kupiqu/fig2svg). Rat brain anatomy graphics were derived from “Brain Maps 4.0” (Larry Swanson; University of Southern California) (Swanson, 2018).

Based on the observation that SA latency data distributions (**Fig. 2E,F**) exhibit characteristics of an exponential distribution, we determined that a non-parametric statistical approach was most appropriate for the behavioral component of this study. Moreover, non-parametric statistical tests were used for electrophysiology data after various tests (Anderson-Darling, D’Agostino & Pearson, Shapiro-Wilk) did not indicate the data were likely to have come from a normal distribution.

## RESULTS

### Rats Self-Administer Nicotine on an Intermittent Access Schedule

Male Sprague Dawley rats (n = 40) were implanted with indwelling jugular catheters and trained to nose poke for intravenous nicotine infusions (0.03 mg/kg/infusion) 2 h/session, 10 sessions (5 sessions per week, Mon-Fri). This continuous access (ContA) phase employed an FR1 schedule of reinforcement (20 s timeout) with visual cues accompanying the nicotine infusion. After ContA, rats were transitioned to intermittent access (IntA) nicotine SA for 7 additional sessions, 4 h/session. IntA involved discrete trials (5 min duration) where nicotine (FR1, 20 s timeout, 0.03 mg/kg/infusion) was available, interspersed with 15 min periods (inter-trial interval; ITI) without drug access. During each ITI, the house light was extinguished and all active/inactive responses were recorded but had no scheduled consequence. During a 4 h IntA session, subjects had 12 discrete trials for a total of 60 min of access to nicotine. See **Fig. 1A** for a schematic of this workflow. Control rats (n = 24) were trained to self-administer saline. Exemplar raster plots show nicotine SA or saline SA training records for ContA (**Fig. 1B,C**) and IntA (**Fig. 1D,E**). Similar to our recent nicotine SA report (Jin et al., 2020), responding on the active nose poke far exceeded responding on the inactive nose poke during ContA of nicotine SA (**Fig. 1F**; session #10 *p* < 0.0001, Mann-Whitney U = 54). This trend was maintained during IntA (**Fig. 1F**; session #17 *p* < 0.0001, Mann-Whitney U = 94.5). Active responding for saline declined during ContA (session #1 vs. #10 *p* < 0.0001, Mann-Whitney U = 89.5) and only transiently increased during the first 1-2 sessions of IntA (**Fig. 1G**). However, active responses for saline did exceed inactive responding (session #17 *p* < 0.0001, Mann-Whitney U = 98.5), likely illustrating the low but measurable reinforcing efficacy of the visual cues associated with nicotine/saline infusions. Consistent with active responding, subjects earned more nicotine infusions vs. saline infusions during ContA (**Fig. 1H**; session #10 *p* < 0.0001, Mann-Whitney U = 130.5). This difference was maintained during IntA (**Fig. 1H**; session #17 *p* < 0.0001, Mann-Whitney U = 95). These results indicate that SD rats reliably acquire nicotine SA and avidly participate in nicotine IVSA under intermittent access conditions.

**Figure 1.**
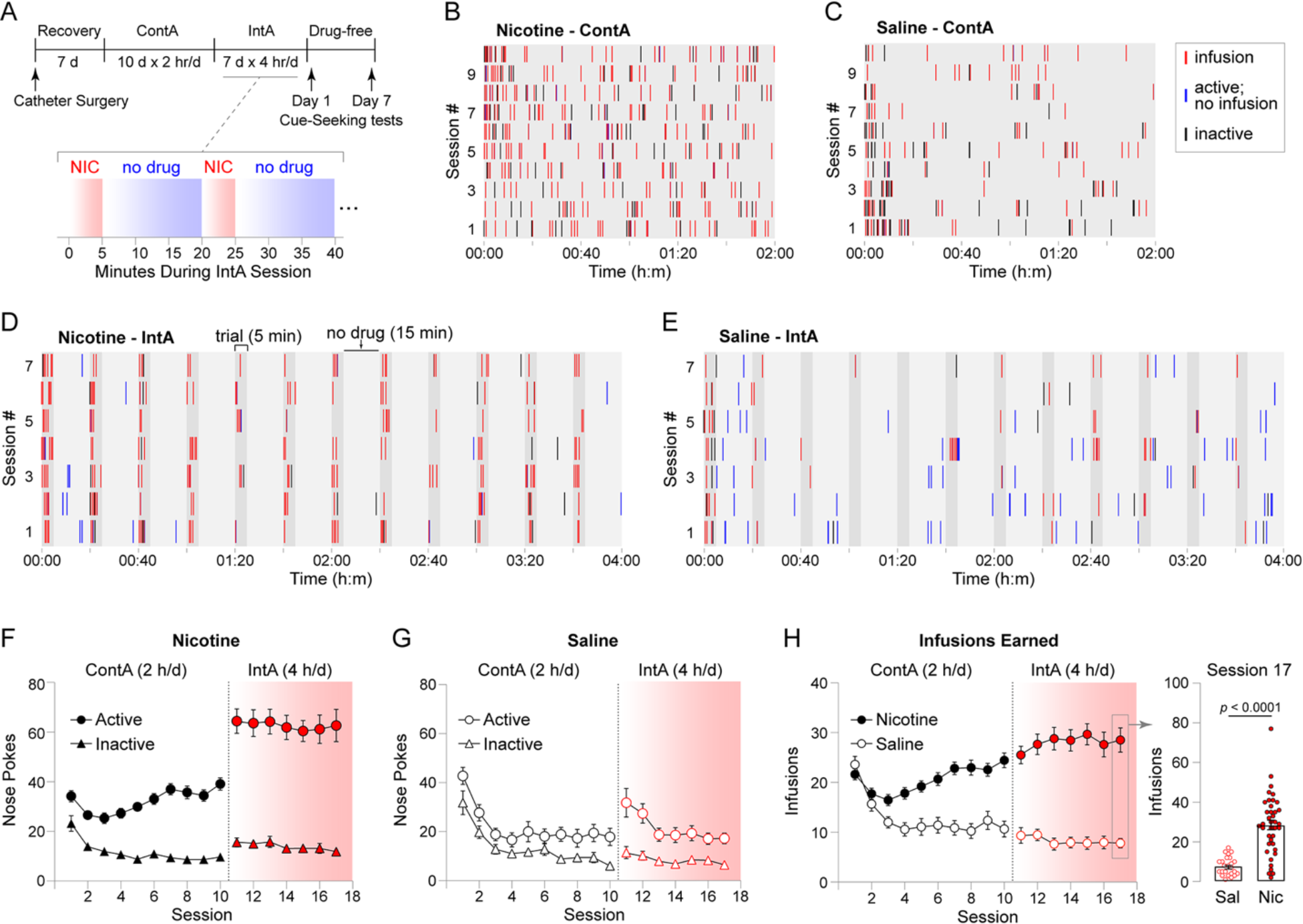
Intermittent access nicotine self-administration. **A**. Self- administration workflow. Rats (nicotine SA, n = 40; saline SA, n = 24) were catheterized and recovered from surgery for 7 d. Rats were trained to self-administer nicotine on an FR1 schedule, 2 h per session for 10 sessions (continuous access; ContA). Next, nicotine access switched to IntA for 7 sessions, 4 h per session; at the start of the session, rats had access to nicotine for 5 min followed by 15 min without access. This cycle repeated 12 times during the 4 h IntA session. After 7 IntA sessions, cue-seeking sessions were conducted on day 1 and day 7 of a drug-free period. **B-C**. Representative 10 d ContA record (infusions, unrewarded active responses, and inactive responses) for a rat trained to respond for nicotine (B) or saline (C). **D-E**. Representative 7 d IntA record for a nicotine SA (D) or saline SA (E) rat. **F-G**. Mean (± SEM) active and inactive responses for n = 40 male nicotine SA rats (F) and n = 24 male saline SA rats (G) are shown for ContA (2h/d) and IntA (4h/d) sessions. **H**. Mean (± SEM) nicotine vs. saline infusions earned are shown for ContA and IntA sessions. At right, a box/dot plot is shown for infusions earned on session #17. *p* value: Mann-Whitney test.

Next, we further explored responding for nicotine during ContA and IntA. We did not analyze data from sessions 1-5 because responding typically did not stabilize until session 6. Infusion times during ContA session #6-10 for n = 36 rats were pooled and expressed as a histogram with 1 min bins (**Fig. 2A**). This revealed that nicotine infusions were more likely during the first ∼10 min of a 2 h ContA session. Subsequently, infusion probability declined and remained stable for the remainder of the session. When the same analysis was conducted on the infusion time data during the 7 IntA sessions, we did not observe the same type of distribution (**Fig. 2B**). Infusion probability was greatest during the first discrete trial, but the likelihood of an infusion did not appear to decline as sharply during subsequent trials during the 4 h IntA session. We further dissected the IntA infusion time data to reveal the probability of an infusion during each minute of the 5 min discrete trial. This indicated an approximately equal probability of an infusion during the first 2 minutes of a nicotine discrete trial, with probability slowly falling during minutes 3, 4, and 5 (**Fig. 2C**).

**Figure 2.**
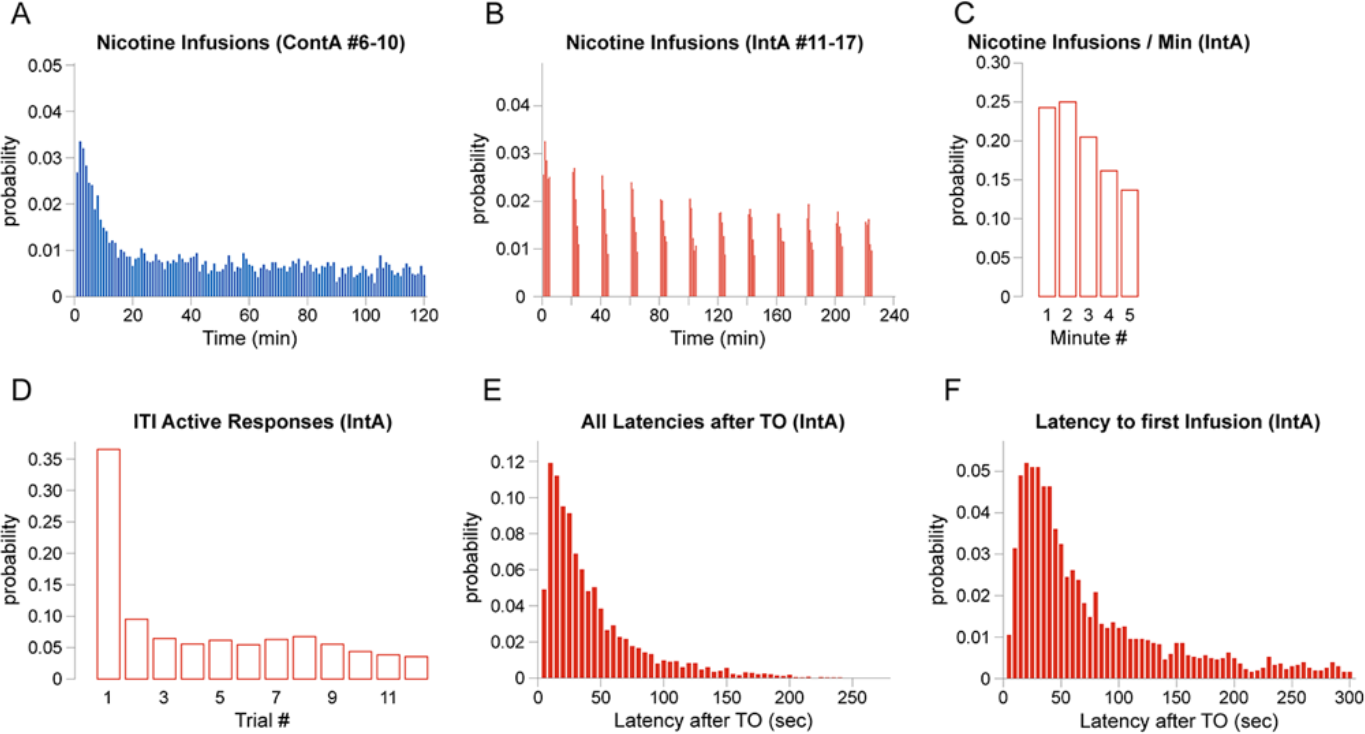
Probabilistic analysis of nicotine self-administration behavior. **A**. Nicotine infusion histogram for sessions 6-10 of ContA. Infusion time data for sessions 6- 10 of nicotine SA ContA pooled (n = 36 rats), binned (width = 1 min), and expressed as a probability histogram to show when nicotine infusions occurred during the later sessions of nicotine SA ContA. **B**. Nicotine infusion histogram for all IntA sessions. Infusion time data for all 7 sessions of nicotine IntA SA pooled (n = 36 rats), binned (width = 1 min), and expressed as a probability histogram to show when nicotine infusions occurred during IntA nicotine SA ContA. **C**. IntA infusions during nicotine access periods. The infusion probability during minute 1, 2, 3, 4, and 5 of nicotine access during IntA sessions are shown for all (n = 36) rats. **D**. Active responses during IntA inter-trial intervals. The probability of an active nose poke during each of 12 intertrial intervals (ITI) are shown. **E.** Infusion latency after timeout. Values for the latency to self-administer a nicotine infusion (time durations) after a timeout period are expressed as a probability histogram. **F.** Infusion latency: first infusion of discrete trial. Values for the latency to self-administer a nicotine infusion after the start of a discrete trial during IntA are expressed as a probability histogram.

To assess how well the rats trained on nicotine SA IntA and learned to disregard the active nose poke during the ITI of an IntA session, we examined the distribution of active responses during each of the 12 ITIs during IntA. The probability of an active response during the first ITI was ∼0.35 but declined dramatically in subsequent ITIs (**Fig. 2D**).

This suggested that rats quickly learned to avoid active responding when nicotine was not available. We saw little to no evidence for “perseverative” responding during the ITI, which has been noted during cocaine SA (Bock et al., 2013). During IntA, a 20 s timeout occurred after each rewarded active response (i.e., each infusion) that resulted in the house light extinguishing for 20 s. After the timeout, the house light was illuminated to signal that nicotine was again available. The time between house light re-illumination and a subsequent rewarded response represents the IntA infusion latency, which is related to urgency or motivation to initiate drug taking (Green et al., 2015; Maccioni et al., 2015). We analyzed the distribution of infusion latencies by binning the latency probability values. The most common latency after a timeout was between 5-10 s, and the distribution of latency times exhibited characteristics of an exponential distribution (**Fig. 2E**). Similarly, we analyzed the distribution of times between the start of a discrete nicotine trial to the first infusion (latency to the first infusion). The shape of the distribution was similar to **Fig. 2E**, with the most common latency to the first infusion being 15-20 s (**Fig. 2F**).

Daily cigarette smokers consume more cigarettes in the morning (Shiffman et al., 2014), consistent with the idea that they “load” on nicotine and then subsequently titrate their consumption throughout the rest of the day to maintain an optimal intake rate. We speculated that ContA nicotine SA would allow for rapid nicotine loading whereas IntA nicotine SA may not. To test this, we modeled plasma nicotine concentrations during ContA and subsequent IntA nicotine SA sessions using a two-compartment pharmacokinetic model. On the 10^th^ session of ContA (when animals are well-trained), modeled plasma nicotine levels reached their maximal level at an average of 84 ± 5 min, with subsequent infusions predicted to maintain plasma nicotine levels at ∼50 ng/mL for the duration of the session (**Fig. 3A**). We also modeled plasma nicotine levels after the same rats were transitioned to IntA nicotine SA. On the 7^th^ IntA session (session #17 overall), modeled plasma nicotine levels peaked at 152 ± 12 min, which was significantly different from time to peak for ContA (**Fig. 3A,B**; *p* < 0.0001, Mann-Whitney U = 292). There was no significant difference between the estimated peak plasma nicotine concentration that was achieved on session #10 of ContA vs. session #7 of IntA (**Fig. 3C**; ContA: 61 ± 4 ng/mL, IntA: 53 ± 4 ng/mL; *p* = 0.207, Mann-Whitney U = 535). To examine nicotine exposure over time under ContA and IntA conditions, we measured area under the curve (AUC) for the estimated plasma nicotine vs. time data after 2 h, 4 h, and 24 h. AUC2h was significantly higher for ContA vs. IntA (**Fig. 3D**; *p* = 0.0002, Mann-Whitney U = 328), but there was no significant difference between ContA and IntA for AUC4h (*p* = 0.699, Mann-Whitney U = 613) and AUC24h (*p* = 0.149, Mann-Whitney U = 519) (**Fig. 3D**).

**Figure 3.**
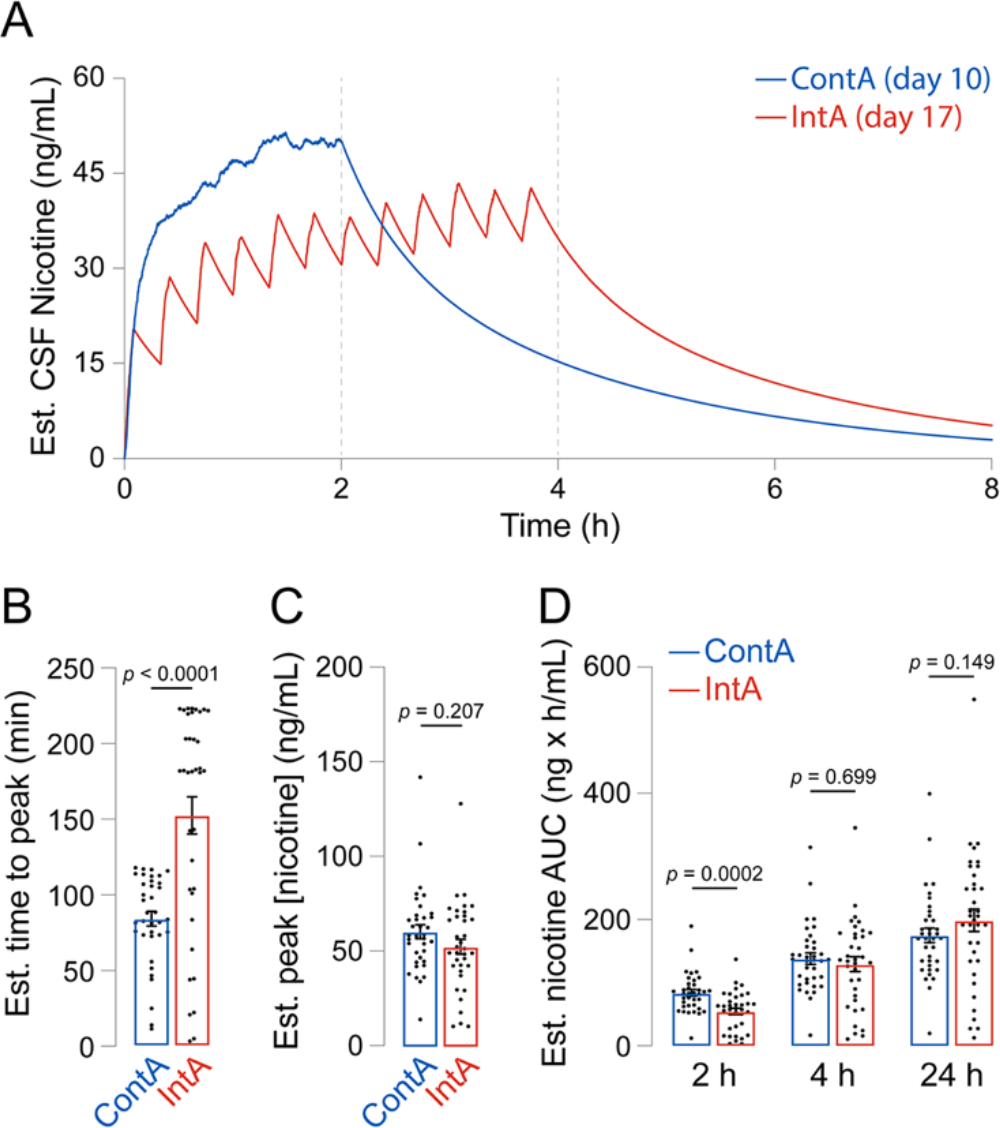
Estimated nicotine pharmacokinetics during SA. A. Estimated plasma nicotine during ContA and IntA nicotine SA. Nicotine infusion vs. time data from Fig. 2 (nicotine SA, n = 36; saline SA, n = 24) was used to model plasma nicotine levels during ContA and IntA SA sessions. Estimated plasma nicotine vs. time on session #10 (ContA) and #17 (IntA) was derived for all nicotine SA rats, and the mean plasma nicotine vs. time profile is shown. **B-C**. Estimated peak plasma nicotine and time to reach peak. For each rat’s nicotine vs. time profile (ContA session #10 and IntA session #17), the time when plasma nicotine level peaked (B) and the peak itself (C) was derived and is shown in the bar/dot plot. *p* value: Mann-Whitney test. **D**. Plasma nicotine exposure over time. The area under the plasma nicotine vs. time curve (AUC) was calculated after 2h, 4h, and 24 h for each rat on session #10 (ContA) and session #17 (IntA). *p* value: Mann-Whitney test.

### Relapse-like behavior after IntA nicotine SA

After nicotine or saline SA acquisition and IntA sessions, we examined relapse-like behavior. During day 1 and day 7 of a drug-free period, rats were presented with the nicotine-associated cues and were allowed to respond on the active or inactive nose poke. During these sessions, nose poke responding activated the nicotine-associated visual cues but did not result in an infusion. A representative cumulative response record for a nicotine SA rat is shown (**Fig. 4A**), indicating that responding on the active nose poke was enhanced on drug-free day 7 compared to day 1. Responding on the inactive nose poke did not change during the drug-free period (**Fig. 4A**). Summary within-subjects results indicate that unreinforced responding was enhanced during the drug-free period (**Fig. 4B**; Active: *p* = 0.01, Wilcoxon W = 56; Inactive: *p* = 0.134, Wilcoxon W = 39). Active responding in the saline SA group was not significantly different on day 7 vs. day 1 (**Fig. 4C**; *p* = 0.094, Wilcoxon W = 17). A difference in inactive responding was noted on day 7 vs. day 1 (**Fig. 4C**; *p* = 0.016, Wilcoxon W = 28). Active response records from all relapse sessions (all examined nicotine SA rats) on day 1 and day 7 are offered as a visual summary measure (**Fig. 4D**). As described for IntA (**Fig. 2F**), we analyzed the distribution of response latencies (latency to respond after TO) during drug-free day 1 and day 7 relapse sessions. There was a significant shift toward shorter latency values in the nicotine SA group on drug-free day 7 compared to day 1 (**Fig. 4E**; *p* = 0.017, Kolmogorov-Smirnov D = 0.1705). We found no such shift in the latency distribution (*p* = 0.245, Kolmogorov-Smirnov D = 0.2014) in the saline SA group (**Fig. 4F**). Mean latency (± SEM) values are plotted for reference (Inset, **Fig 4E,F**).

**Figure 4.**
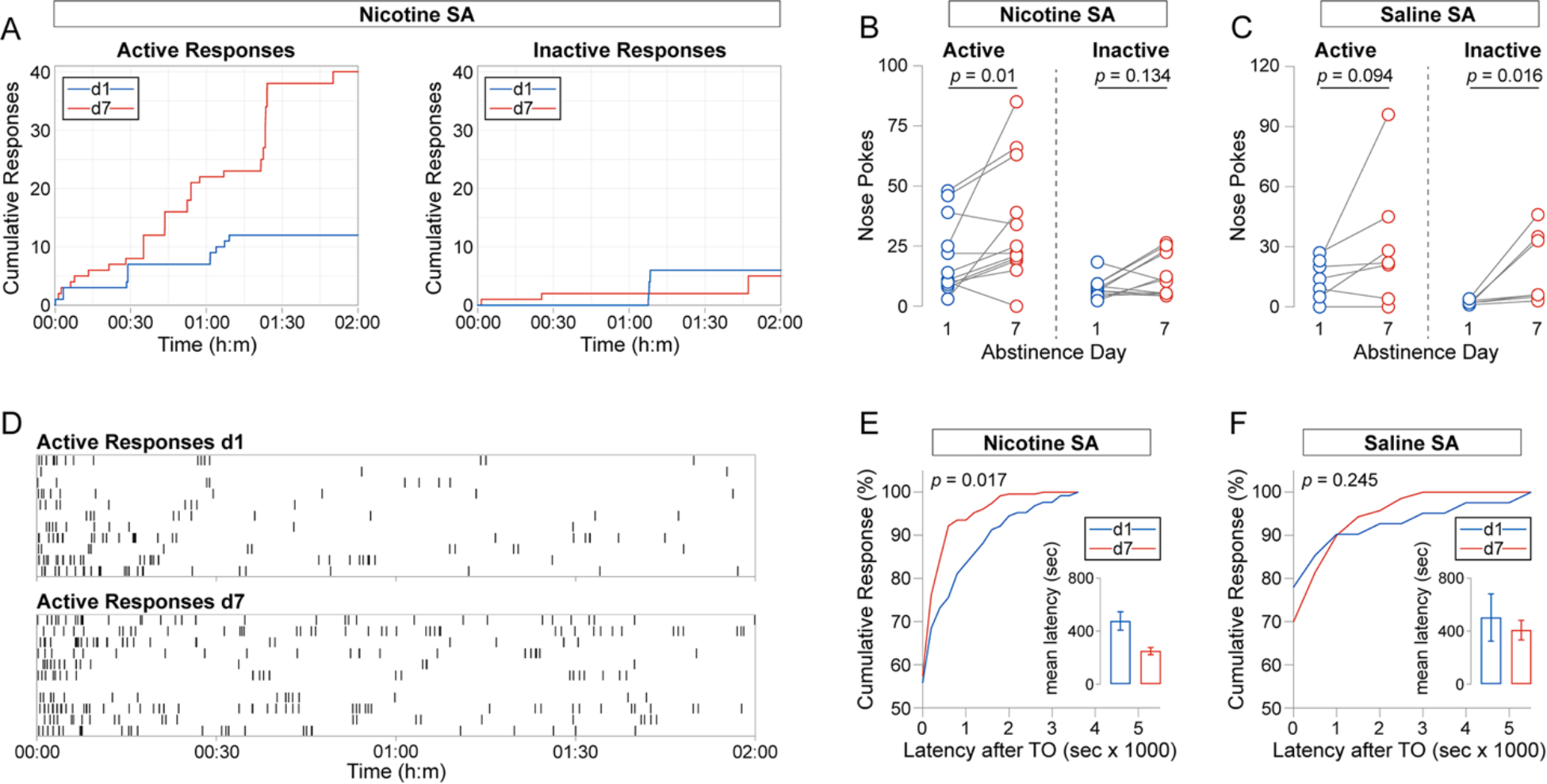
Relapse-like behavior after IntA nicotine SA. **A**. Cumulative responses during nicotine-seeking sessions. Representative cumulative active and inactive response records are shown for a nicotine SA rat on nicotine seeking session d1 and d7. **B-C**. Summary of nicotine-seeking behavior. Before-after plots are shown for active and inactive responses on d1 and d7 in the nicotine SA (B; n = 12) and saline SA (C; n = 7) group. *p* value: Wilcoxon matched-pairs signed rank test. **D**. Nicotine-seeking on d1 vs. d7; visual summary. A raster plot is shown for all nicotine SA animals on d1 and d7. Vertical tick marks denote active responses; each row shows a specific subject’s response record. **E-F**. Cumulative distribution of active response latencies (latency to respond after a TO) on d1 and d7 for all nicotine SA (E) and saline SA (F) rats. Inset: mean latency values on d1 and d7. *p* value: Kolmogorov-Smirnov test.

### Neurobiological changes associated with IntA nicotine SA

If these cue-induced nicotine seeking data (**Fig. 4**) are biologically meaningful, we reasoned that there should be some corresponding neurobiological changes in brain regions that are known to play a role in nicotine relapse. To address this hypothesis, we performed quantitative bulk proteomic analysis of dorsal striatal extracts from rats after saline or nicotine SA (10 ContA sessions and 7 IntA sessions, as described above) and 1 or 7 drug-free days. We used tandem mass tag (TMT) reagents enabling the identification and relative multiplexed quantitation of the same set of peptides and proteins in complex mixtures across all biological replicates (Jongkamonwiwat et al., 2020; Rao and Savas, 2021). We selected dorsal striatum based on 1) prior proteomic studies that examined dopamine system brain tissue following nicotine (Lee et al., 2021) or cocaine (Shen et al., 2016) treatment, 2) reports suggesting that chronic nicotine treatment alters the physiology of neurons in the dorsal striatum (Xia et al., 2017), and 3) its role in relapse-like behavior (Gerdeman et al., 2003; Everitt and Robbins, 2016). After SA and 1 or 7 drug-free days, the dorsal striatum was dissected and proteins were extracted, digested to peptides, labeled with an individual TMT tag, and pooled (**Fig. 5A**). The labeled peptides were fractionated and analyzed with the MS3 multinotch method [33, 34]. We quantified 4,399 protein groups across the 16 samples. When we compared the four groups, we found significantly altered proteins in the comparison of day 7 saline vs. day 7 nicotine groups. To assess the reliability of these measures, we performed hierarchical clustering of the 220 significantly altered proteins (FDR < 0.05) and found the biological replicates within the groups are highly similar (**Fig. 5B**). To visualize the complete proteomic dataset, we plotted the fold-change (FC) and *p*-value for all the quantified proteins and found that a similar number of proteins had elevated and reduced FC (**Fig. 5C**, Extended Data **Fig. 5-1**). To further assess protein FC, we generated a heat map of the 220 significantly altered proteins (FDR < 0.05). This illustrates the trends across the individual biological replicates (**Fig. 5D**). Next, to investigate the possibility that proteins with common functions are co-regulated during the drug-free days, we subjected the proteins with significantly (FDR < 0.05) elevated (n=91) or reduced (n=129) FC to Gene Ontology (GO) molecular function enrichment analysis. Notably, only the pool proteins with elevated levels were significantly enriched for common functions. These functions were predominantly related to membrane signaling and the activation of gene transcription (**Fig. 5E**).

**Figure 5.**
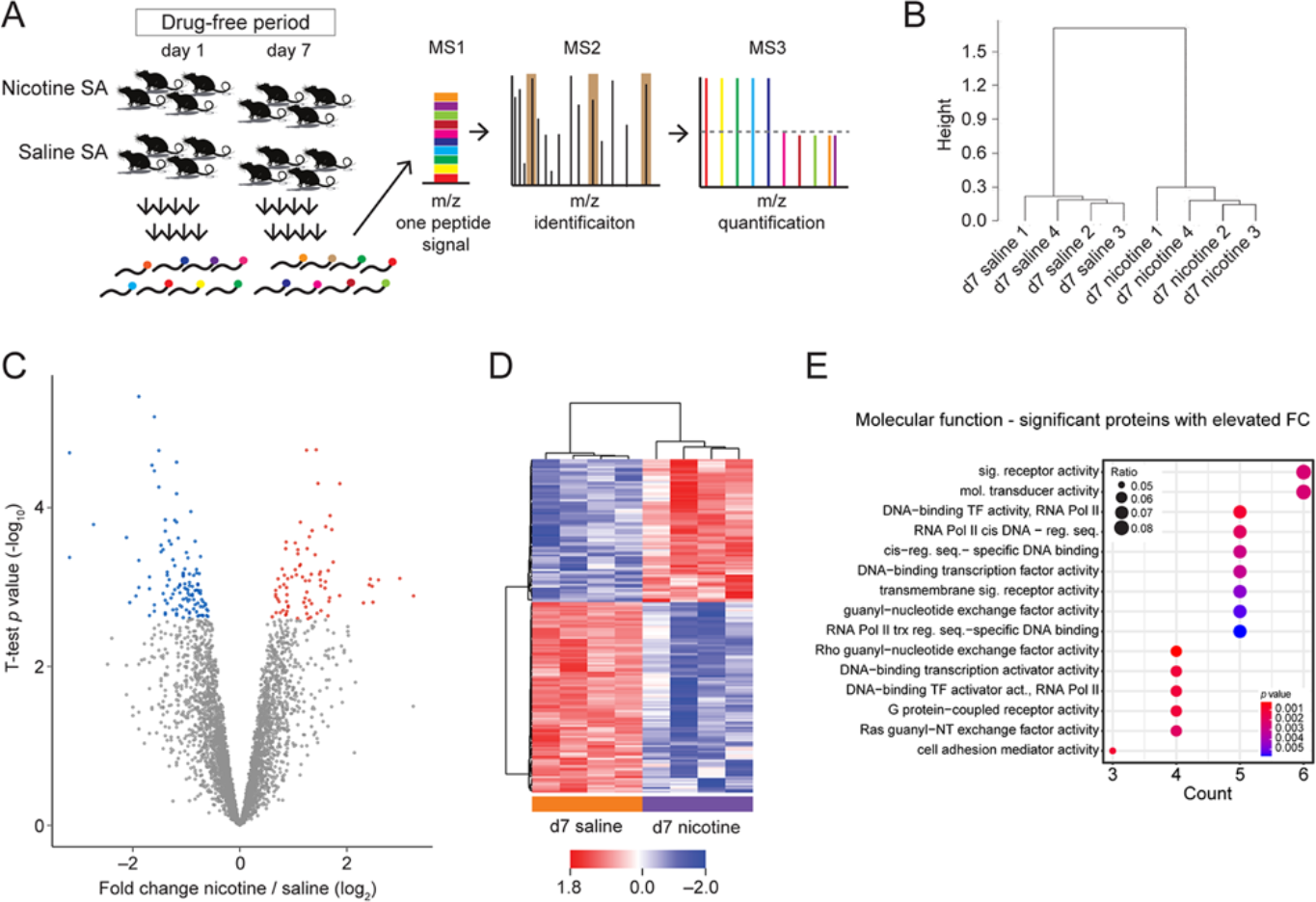
Identification of dorsal striatal proteins with altered levels after IntA nicotine SA. **A**. Experimental workflow to identify proteins with altered fold change (FC) in the dorsal striatum during incubation of craving. Proteins were extracted from the dorsal striatum (n = 16 male rats, n = 4 per group, 4 indicated groups), digested to peptides, labeled with individual TMT tags, and mixed in equal ratios. The peptides were identified and quantified with LC-MS3-based proteomics. **B**. Hierarchical clustering dendrogram of the individual biological replicates/datasets based on the 220 significantly altered proteins in the nicotine and saline groups on drug-free day 7 (FDR < 5%). **C**. Volcano plot depicting the FC (i.e. nicotine / saline) and *t*-test *p*-value. N = 4399 protein groups were quantified. Proteins with significantly elevated (n = 93) or reduced (n = 129) FC and FDR < 5% are colored in red or blue, respectively. **D**. Heat map depicting protein FCs for significant proteins (FDR < 5%) from the day 7 saline and nicotine groups. **E**. Summary results from gene Ontology (GO) enrichment analysis (molecular function) from the 91 proteins with significantly elevated FC. The statistics for the GO over-representation analyses were performed using hypergeometric *p*-values. Details of elevated and reduced proteins are included in Extended Data Figure 5-1.

To further explore the neurobiological changes induced by IntA nicotine SA, we measured [^11^C]nicotine radioactive whole-brain uptake in IntA nicotine (and saline control) SA rats with micro positron emission tomography (PET). This approach has been used successfully in human subjects to examine nicotine uptake in cigarette and electronic cigarette smokers (Solingapuram Sai et al., 2020). Rats were trained as indicated in **Fig. 1A**, and [^11^C]nicotine radioactive uptake in the brain was measured as standard uptake values (SUV in KBqML*ccm) on the day after the 7^th^ IntA session. An initial PET scan indicated that nicotine SA rats exhibited ∼27% greater [^11^C]nicotine uptake compared to saline SA controls (**Fig. 6A**). This prompted us to attempt to independently replicate these results. The replication experiment yielded consistent results (∼40% increase in brain radioactive uptake in nicotine SA vs. saline SA). These data were pooled based on this consistency, yielding a significant increase in [^11^C]nicotine uptake in nicotine SA vs. saline SA rats (nicotine SA SUV: 46.1 ± 3.5, n = 4; saline SA SUV: 34.4 ± 1.2, n = 6; *p* = 0.019, Mann-Whitney U = 1) (**Fig. 6B**). To further validate the *in vivo* imaging results of the microPET study, brains were extracted after the scans, and radioactivity was measured using a γ-counter. The residual radioactivity (‘percent injected dose per gram (%ID/g)’) was measured, revealing a pattern consistent with the PET imaging: nicotine SA brain radioactivity (%ID/g = 3.58 ± 0.6) was ∼97% greater compared to saline SA (%ID/g = 1.81 ± 0.2).

**Figure 6.**
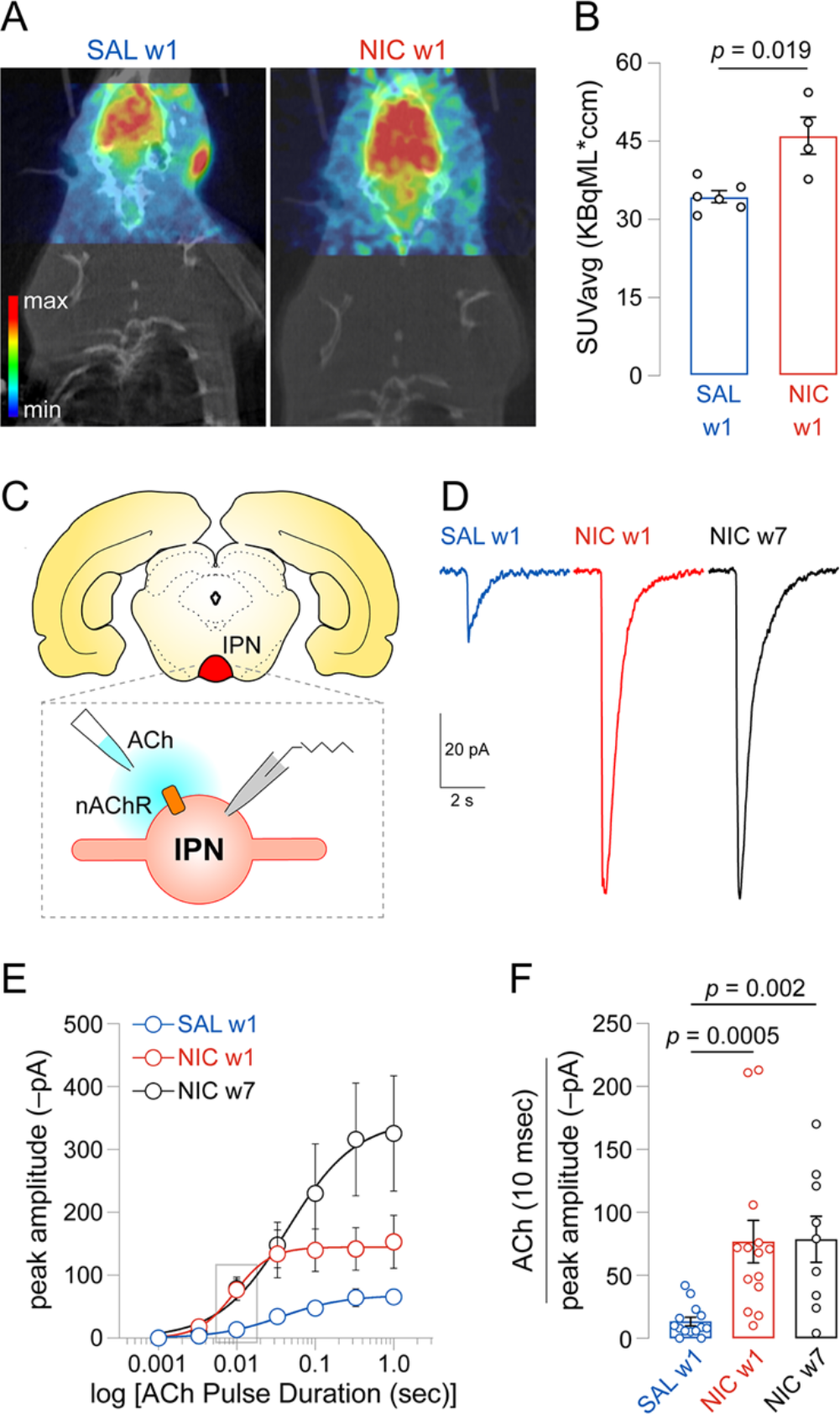
nAChR upregulation after IntA nicotine SA. **A**. Example PET imaging micrograph for saline SA or nicotine SA rats injected with [^11^C]nicotine radiotracer on withdrawal day 1 (w1). **B**. Summary [^11^C]nicotine uptake data for saline SA (n = 6) and nicotine SA (n = 4) rats on day w1. *p* value: Mann-Whitney test. **C**. Experiment schematic. A rat coronal brain section containing the IPN is shown. Inset: IPN neurons are recorded via whole-cell patch-clamp electrophysiology and nAChRs are activated by local application of ACh. **D**. nAChR responses. Slices were prepared from saline SA rats on withdrawal d1 (n = 3), and slices were prepared from nicotine SA rats on withdrawal d1 (n = 5) or d7 (n = 4); no seeking sessions were run. All ACh-elicited responses were averaged and the average trace is shown for pulse duration = 33 ms. **E-F**. Summary data for nAChR sensitization after IntA. For each cell, multiple responses were recorded at different ACh pulse durations, as indicated. A scatter plot (for all cells) is shown (F) for 10 ms data. *p* value: Dunn’s multiple comparison test adjusted *p* value after omnibus Kruskal-Wallis test (Kruskal Wallis statistic = 17.11, *p* = 0.0002).

To complement the micro-PET results and to further examine nAChR upregulation with functional measurements, we examined nAChR functional activity with patch clamp electrophysiology after rats were trained on IntA nicotine SA. The interpeduncular nucleus (IPN), which directly modulates the serotonergic raphe nuclei (Quina et al., 2017; Morton et al., 2018) and indirectly modulates the mesolimbic dopamine system (Wolfman et al., 2018), plays an important role in regulating nicotine intake (Fowler et al., 2011). Although we previously demonstrated sensitized functional nicotinic acetylcholine receptors in IPN following chronic passive nicotine exposure in mice (Arvin et al., 2019), whether/how cholinergic signaling is sensitized by nicotine SA is unknown.

After training rats to self-administer nicotine (see **Fig. 1A**), brain slices were prepared and whole cell patch clamp recordings were made from neurons in the rostral IPN (IPR) (**Fig. 6C**). This brain area was chosen due to its high level of nAChR functional activity (Arvin et al., 2019). ACh was locally applied to the recorded cell to activate nAChRs and measure their level of functional activity. Rats from saline SA (n = 3) and nicotine SA (n = 5) groups were studied on day 1 after the last IntA session, denoted withdrawal day 1 (w1). We also examined IPN nAChRs in the nicotine SA group on withdrawal day 7 (w7; n = 4). Rats destined for brain slice patch clamp physiology experiments were treated according to the workflow in **Fig. 1A**, except that no relapse sessions were conducted. Compared to nAChR responses from saline SA w1 IPN neurons, responses from nicotine SA w1 IPN neurons were of greater amplitude (**Fig. 6D**). Intriguingly, nAChR responses in IPN neurons remained elevated after 7 days of withdrawal in the nicotine SA group (**Fig. 6D**). Measuring ACh responses across a range of ACh pulse durations (**Fig. 6E**), which approximates a concentration-response curve, more readily demonstrates nAChR functional sensitization following IntA nicotine SA. At an ACh pulse duration of 10 ms (**Fig. 6F**), we noted an overall effect of treatment condition (*p* = 0.0002, Kruskal-Wallis statistic = 17.11), with significant differences between saline SA w1 vs. nicotine SA w1 (adjusted *p* = 0.0005) as well as saline SA w1 vs. nicotine SA w7 (adjusted *p* = 0.002) using Dunn’s multiple comparison testing.

### Behavioral mechanisms of IntA nicotine SA

Next, we compared nicotine intake and estimated pharmacokinetics in the intermittent vs. continuous access paradigms. Rats were trained on ContA nicotine SA for 10 sessions and were then transitioned to one of the following conditions for sessions 11-17: 1) IntA 4 h, 2) ContA 2h, or 3) ContA 4h. The IntA 4h condition is identical to the previous IntA experiments described in this study.

Rats were tested for cue-induced nicotine seeking on days 1 and 7 of a nicotine-free period, which began on the day after SA session #17 (**Fig. 7A**). The 3 groups acquired nicotine SA in a similar manner (**Fig. 7B**); there was no significant difference between total infusions during day 1-10 (acquisition) in the ContA 2h or ContA 4h group compared to IntA 4h (ContA 2h: *p* = 0.34, Mann-Whitney U = 41; ContA 4h: *p* = 0.71, Mann-Whitney U = 59.5). During the ensuing 7-session test period, infusions were marginally increased in the IntA group compared to ContA 2h (*p* = 0.041, Mann-Whitney U = 26) and there was a non-significant trend toward increased infusions compared to the ContA 4h group (*p* = 0.085, Mann-Whitney U = 34) (**Fig. 7C**). These results are remarkable given that rats in the IntA paradigm have two-fold (vs. ContA 2h) or four-fold (vs. ContA 4h) less time to access nicotine. We modeled plasma nicotine levels in these groups based on their SA infusion time records, and the average plasma nicotine vs. time profile is shown for session #17 (**Fig. 7D**). Relative to the IntA group, ContA 2h and ContA 4h rats reached their peak nicotine levels sooner (vs. ContA 2h: *p* = 0.0001, Mann-Whitney U = 5; vs. ContA 4h: *p* = 0.0005, Mann-Whitney U = 13) (**Fig. 7E**). Whereas ContA 2h rats achieved a similar estimated peak plasma nicotine level compared to IntA rats (*p* = 0.282, Mann-Whitney U = 39), peak plasma nicotine levels in ContA 4h rats was significantly lower than in IntA rats (*p* = 0.002, Mann-Whitney U = 17) (**Fig. 7F**). Similar to our within-subject analysis (**Fig. 3D**), AUC2h was greater for the ContA 2h group (*p* = 0.005, Mann-Whitney U = 16) whereas AUC4h (*p* = 0.918, Mann-Whitney U = 53) and AUC24h (*p* = 0.197, Mann-Whitney U = 36) did not differ significantly (**Fig. 7G**). In the ContA 4h group, AUC2h was similar to IntA levels (*p* = 0.413, Mann-Whitney U = 52). In ContA 4h there was a trend toward reduced AUC4h (*p* = 0.051, Mann-Whitney U = 34) compared to IntA, and AUC24h (*p* = 0.023, Mann-Whitney U = 29) was significantly lower (**Fig. 7G**).

**Figure 7.**
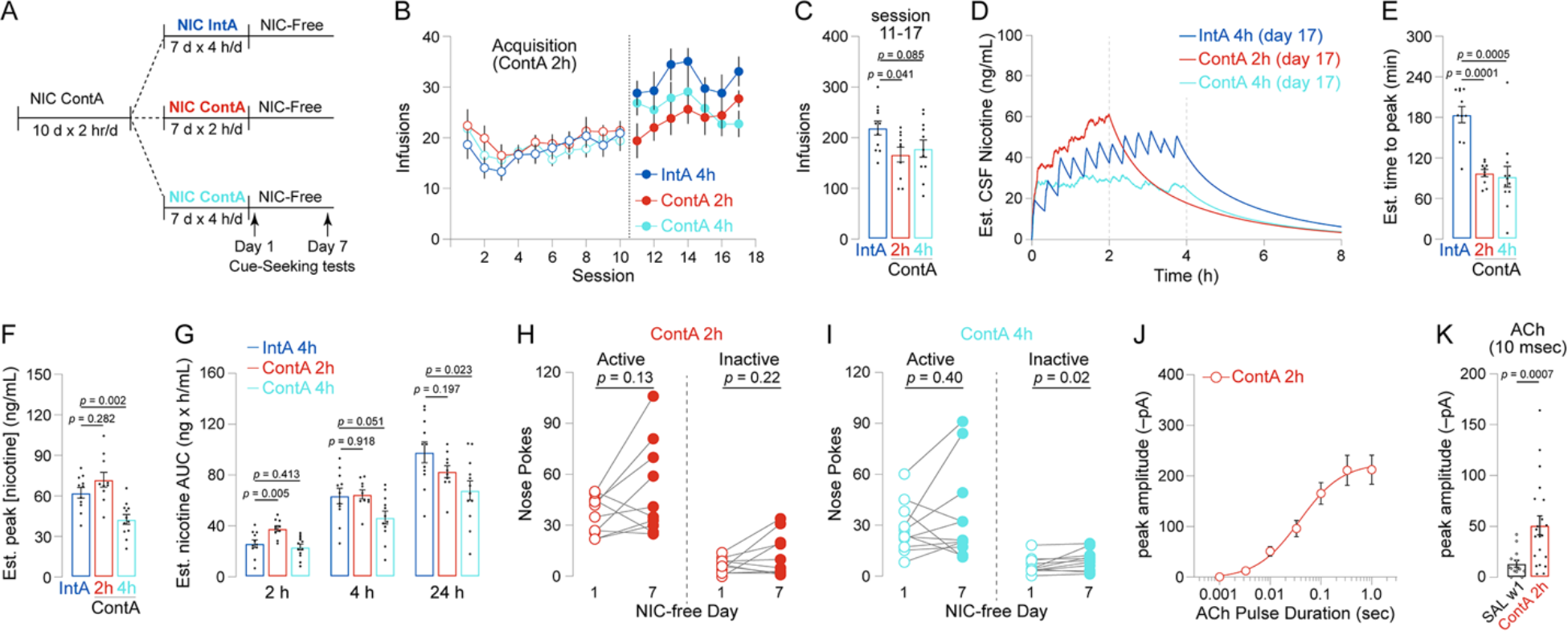
ContA vs IntA nicotine SA. **A**. Experimental schematic. Rats (n = 33) were trained on the identical ContA paradigm for 10 sessions and were then assigned to one of three groups: IntA (n = 11 were transitioned to IntA as described in Fig. 1A), ContA 2h (n = 10 continued on ContA 2 h/d for another 7 sessions), or ContA 4h (n = 12 were transitioned to ContA 4 h/d for another 7 sessions). Cue-seeking tests were conducted on day 1 and 7 of a subsequent nicotine-free period. **B**. Nicotine infusions during sessions 1- 17 are shown for the indicated groups. **C**. Summary dot/bar plot shows the total number of infusions for each rat for sessions 11-17. *p* value: Mann-Whitney test. **D**. Estimated nicotine pharmacokinetics for the indicated ContA (2h and 4h) and IntA groups. Infusion data for each rat was used to model the plasma nicotine vs. time profile on the last day (session #17) of training, and an average profile was derived and plotted. **E-G**. Using the data shown in (D), the estimated time to reach peak plasma [nicotine] (E), the peak plasma [nicotine] (F), and the area under the curve at 2 h, 4 h, and 24 h (G) is shown. *p* value: Mann-Whitney test. **H-I**. Summary of nicotine-seeking behavior for the ContA 2h (H) and ContA 4h (I) groups. Before-after plots are shown for active and inactive responses on d1 and d7 in the indicated ContA group. *p* value: Wilcoxon matched-pairs signed rank test. **J-K**. Summary data for IPN nAChR activity after ContA 2h nicotine SA. For each cell, multiple responses were recorded at different ACh pulse durations, as indicated (J). A scatter plot (for all cells) is shown (K) for 10 ms data. *p* value: Mann- Whitney test with ContA 2h data compared to the control SAL w1 group shown in Fig. 6F.

ContA 2h nicotine SA rats did not exhibit incubation of craving (**Fig. 7H**; active: *p* = 0.13, Wilcoxon W = 31; inactive *p* = 0.22, Wilcoxon W = 21). Likewise, ContA 4h nicotine SA rats did not exhibit incubation (**Fig. 7I**; active: *p* = 0.40, Wilcoxon W = 20; inactive *p* = 0.02, Wilcoxon W = 38). Finally, we examined nAChR functional upregulation in IPN neurons of ContA 2h rats. As with our standard IntA treatment (**Fig. 1A**), 17 sessions of ContA nicotine SA led to enhanced ACh-evoked currents in neurons of the dorsal IPN (**Fig. 7J,K**; *p* = 0.0007, Mann-Whitney U = 50).

Previous work with cocaine IntA SA (Morgan et al., 2002; Zimmer et al., 2011; Calipari et al., 2013), upon which this study was modeled, did not utilize a timeout or an associated house light during the 5 min access periods of IntA. To determine whether our inclusion of a timeout (with the associated visual cues) impacted nicotine IntA SA, we trained a new group of rats (n = 11) on a modified IntA paradigm that did not include a timeout (or extinguishment of the house light) during IntA (IntA ‘no TO’; **Fig. 8A**). We hypothesized that removal of the timeout would enhance nicotine intake because total access time would be increased compared to IntA with a 20 s timeout. We compared results from the IntA ‘noTO’ rats to the subset of our initial IntA subjects that completed relapse sessions on day 1 and day 7. These are the same control rats whose data are shown in **Fig. 7**. Contrary to our hypothesis, the lack of a TO and associated visual cues reduced infusions during days 11-17 compared to IntA rats (**Fig. 8B,C**; *p* = 0.038, Mann-Whitney U = 29). We modeled plasma nicotine pharmacokinetics using infusion data from session #17, which revealed a blunted response in IntA ‘no TO’ rats compared to IntA rats (**Fig. 8D**). This reduction in plasma nicotine levels is likely explained by a significant increase in the number of 5-min trials without an active response (omitted trials) in the IntA ‘noTO’ group compared to the IntA group (**Fig. 8D, inset**; *p* = 0.002, Mann-Whitney U = 15).

**Figure 8.**
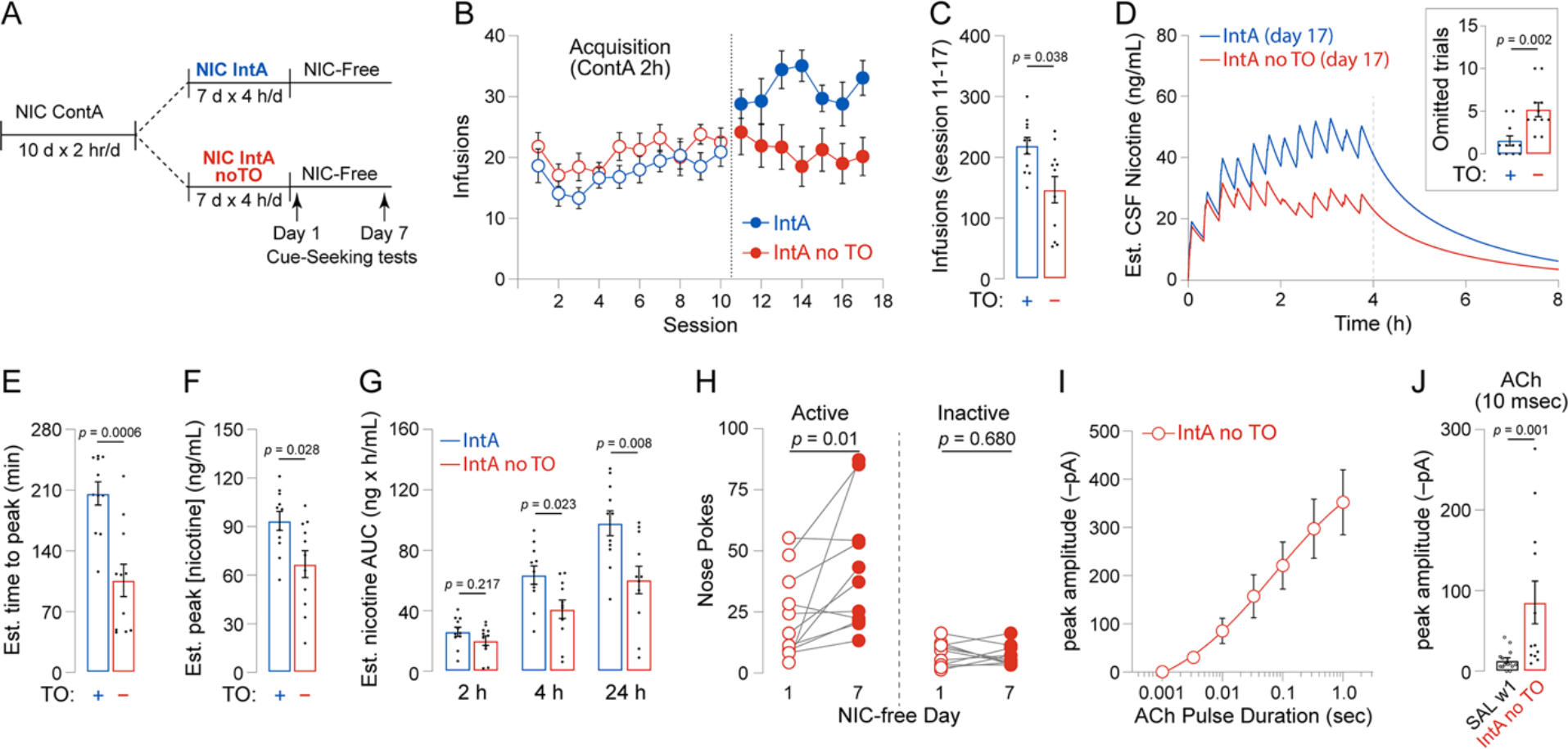
Cue conditions influence nicotine intake during IntA. **A**. Experimental schematic. Rats (n = 22) were trained on the standard ContA paradigm (2 h/d, 10 d, FR1, 0.03 mg/kg/inf) for 10 sessions and were then assigned to one of two groups: IntA (n = 11; 4 h/d, 7 d, FR1, 5 min access, 15 min without access, 0.03 mg/kg/inf) or IntA with no timeout period after an infusion (n = 11). Cue-seeking tests were conducted on day 1 and 7 of a subsequent nicotine-free period. The IntA group was previously shown in Fig. 7 and is shown again here for reference. **B**. Nicotine infusions for the IntA and IntA ‘no TO’ groups are shown. **C**. Summary dot/bar plot shows the total number of infusions for each rat for sessions 11-17. *p* value: Mann-Whitney test. **D**. Estimated nicotine pharmacokinetics for the IntA and IntA ‘no TO’ groups. Infusion data for each rat was used to model the plasma nicotine vs. time profile on the last day (session #17) of training, and an average profile was derived and plotted. Inset: number of omitted trials on session #17 for IntA and IntA ‘no TO’ groups. **E-G**. Using the data shown in (D), the estimated time to reach peak plasma [nicotine] (E), the peak plasma [nicotine] (F), and the area under the curve at 2 h, 4 h, and 24 h (G) is shown. *p* value: Mann-Whitney test. **H**. Summary of nicotine-seeking behavior for the IntA ‘no TO’ group. Before-after plots are shown for active and inactive responses on d1 and d7 in the IntA ‘no TO’ group. *p* value: Wilcoxon matched-pairs signed rank test. **I-J**. Summary data for IPN nAChR activity after IntA ‘no TO’ nicotine SA. For each cell, multiple responses were recorded at different ACh pulse durations, as indicated (I). A scatter plot (for all cells) is shown (J) for 10 ms data. *p* value: Mann-Whitney test with IntA ‘no TO’ data compared to the control SAL w1 group shown in Fig. 6F.

IntA ‘no TO’ rats reached their peak plasma nicotine level earlier in the session than rats that included a TO (**Fig. 8E**; *p* = 0.0006, Mann-Whitney U = 11), but the peak level achieved was lower in the IntA ‘no TO’ group compared to control IntA rats (**Fig. 8F**; *p* = 0.028, Mann-Whitney U = 27). Control IntA (with TO) rats exhibited similar AUC at 2 h (*p* = 0.217, Mann-Whitney U = 41) and greater AUC at 4 h (*p* = 0.023, Mann-Whitney U = 26) and 24 h (*p* = 0.008, Mann-Whitney U = 21) after the start of the session (**Fig. 8G**). Despite this reduction in nicotine exposure in the IntA ‘no TO’ group, incubation of craving was preserved (**Fig. 8H**; active: *p* = 0.01, Wilcoxon W = 55; inactive: *p* = 0.68, Wilcoxon W = −9). Finally, we examined nAChR functional upregulation in IPN neurons of IntA ‘no TO’ rats. As with our standard IntA treatment (**Fig. 1A**), IntA ‘no TO’ nicotine SA led to enhanced ACh-evoked currents in neurons of the dorsal IPN (**Fig. 8I,J**; *p* = 0.0007, Mann-Whitney U = 50).

## DISCUSSION

In this study, we demonstrate that rats will readily self-administer nicotine during 5 min access periods that are interspersed with longer (15 min) periods without nicotine access. This mode of nicotine SA led to enhanced nicotine seeking after a drug-free period (incubation of craving) and similar or greater nicotine intake (yet similar nicotine exposure) compared to ContA SA. Incubation of craving was associated with alterations to the dorsal striatum proteome. IntA nicotine SA is sensitive to the cue conditions, as exclusion of a timeout and the associated visual cues reduced nicotine SA. IntA nicotine SA provoked nAChR upregulation in the interpeduncular nucleus, a brain area involved in nicotine dependence. Accordingly, this study contributes a translationally relevant nicotine exposure paradigm as well as new mechanistic insights into nicotine addiction.

### Nicotine SA: ContA vs. IntA

In our study, rats had only 1 h of access time to take nicotine during a 4 h IntA session compared to 2 h of access during a 2 h ContA session, yet nicotine intake (infusions) was modestly enhanced during IntA (**Fig. 7C**). Pharmacokinetic modeling of plasma nicotine levels provided important insights. ContA (2 h session) SA enabled rats to rapidly increase their plasma nicotine levels within ∼10- 20 min, followed by a phase with slower intake (maintenance) for the duration of the session (**Fig. 3A** and **Fig. 7D**). IntA SA was associated with a qualitatively slower rise in plasma nicotine levels (**Fig. 3A** and **Fig. 7D**). These interpretations are supported by our analysis of infusion probability over time (**Fig. 2A,B**), and are generally consistent with human cigarette smoking behavior during the day (Shiffman et al., 2014). For ContA 2h rats, we found no significant difference between the area under the plasma nicotine vs. time curve at 4 h or 24 h after the start of SA (**Fig. 7G**). For ContA 4h rats, cumulative nicotine exposure over 24 h was reduced compared to IntA (**Fig. 7G**). Although IntA can result in a greater number of infusions than ContA (2 h ContA vs. 4 h IntA; **Fig. 7C**), estimated plasma nicotine pharmacokinetics suggest that when animals self-administer nicotine intermittently, *more infusions are required* to maintain a similar level of nicotine exposure over time. Extrapolating these results to human consumers of combusted tobacco products, an intermittent pattern of intake may lead to similar or even greater exposure to toxic and/or carcinogenic constituents in tobacco smoke.

Rats quickly learned to disregard the nose poke during the ITI of IntA sessions, which was signaled by extinguishment of the house light. Active responses for nicotine were much more probable during the first ITI relative to the rest (**Fig. 2D**). We found no evidence for “perseverative” or “anticipatory” responding for nicotine during the ITI, which has been noted for cocaine when access is temporarily revoked during SA sessions (Bock et al., 2013). We were initially surprised that exclusion of the timeout and the associated visual cues was sufficient to significantly reduce IntA nicotine SA. However, environmental cues play a very important role in animal models of nicotine SA and in human tobacco consumption (Conklin et al., 2008; Conklin et al., 2010). For example, early nicotine SA experiments examining the impact of removing the visual cues (but retaining nicotine infusions) from the SA paradigm revealed that such cues become conditioned reinforcers that are of nearly equal importance compared to nicotine itself (Caggiula et al., 2001). A recent preclinical study corroborated these results, showing that cues previously associated with a standard nicotine dose are sufficient to maintain responding for an unrewarding dose of nicotine (Powell et al., 2020). In our ‘no TO’ experiment, a response on the active nose poke illuminated the cue light inside the nose poke but the house light did not cycle off like it normally would in our typical experiments. Given the importance of visual cues in sustaining nicotine SA at maximal levels (Donny et al., 1999; Caggiula et al., 2001; Chaudhri et al., 2005), we speculate that eliminating this light cue may be the main driver of this behavioral result.

### IntA Nicotine SA and Incubation of Craving

Nicotine exposure via IntA SA is sufficient to not only elicit nicotine seeking on the first day of a drug-free period, but it also supports enhanced seeking after 7 drug-free days. This phenomenon, known as incubation of craving, is a feature of human drug addiction (Gawin and Kleber, 1986; Bedi et al., 2011; Wang et al., 2013; Li et al., 2015; Parvaz et al., 2016). Although previous studies have demonstrated incubation of craving after intravenous nicotine SA in rats (Funk et al., 2016; Markou et al., 2018), nicotine incubation of craving has not previously been reported following an IntA paradigm. Excluding a timeout and the associated house light cues did not impact incubation of craving (**Fig. 8H**). However, in our hands, 17 continuous FR1 SA sessions (2 h or 4 h duration) were not associated with a statistically significant increase in seeking on day 7 vs. day 1 of the drug-free period (**Fig. 7I**). This result differs from a recent study that did note incubation of craving with conditions similar to our ContA groups (Markou et al., 2018). This may be due to sampling variation, as there appears to be a trend in our data (**Fig. 7I**) suggestive of incubation in a fraction of animals. If so, the significant increase in unreinforced responding we report for IntA (**Fig. 4**) could reflect a greater ability of intermittent nicotine (vs. continuous nicotine SA) to establish the neurobiological mechanisms giving rise to incubation of craving. This is speculation; future experiments will be needed to test this hypothesis and further probe the difference in incubation of craving we note between ContA and IntA.

Our discovery-based proteomic analysis revealed that the dorsal striatum undergoes robust changes during a drug-free period after IntA nicotine SA. In particular, we detected greater proteome remodeling after nicotine SA and 7 nicotine-free days vs. saline SA and 7 saline-free days. GO analysis revealed that the pool of significantly elevated proteins was selectively enriched for receptors and transcription factors, suggesting that dorsal striatum signal transduction and gene expression may play an important role in the induction of nicotine craving. Although there are no reports detailing proteomic changes associated with nicotine incubation of craving, the types of enriched pathways we discovered are generally consistent with proteomic changes observed in other chronic nicotine paradigms. For example, Picciotto and colleagues noted proteomic changes in dopamine signaling, GABA signaling, and protein synthesis machinery pathways following chronic nicotine exposure in mice (Lee et al., 2021). Moreover, incubation of cocaine craving in rats was recently shown to be associated with robust changes to several signal transduction pathways in the central nucleus of the amygdala (Hamor et al., 2020).

### nAChR Upregulation after IntA

nAChR upregulation is a remarkably robust effect of chronic nicotine exposure; upregulation occurs in cell lines, animal models, and human smokers (Marks et al., 1983; Schwartz and Kellar, 1983; Perry et al., 1999; Kuryatov et al., 2000; Kuryatov et al., 2005). We recently validated the utility of PET imaging using [^11^C]nicotine for studying nAChRs in vivo during nicotine exposure via cigarette smoking and electronic cigarette usage (Solingapuram Sai et al., 2020). Our PET imaging experiments indicate increased [^11^C]nicotine uptake in nicotine SA brain compared to saline SA, which is suggestive of nAChR upregulation. Similarly, Le Foll and colleagues used PET to detect an increase in 2-[^18^F]fluoro-A-85380 binding to α4β2 nAChRs in non-human primates after intravenous nicotine SA (Le Foll et al., 2016). Our results indicate that the [^11^C]nicotine radiotracer is a useful PET imaging ligand for translational studies examining nAChRs during and after nicotine exposure *in vivo*.

We and others previously reported that chronic passive nicotine exposure is sufficient to enhance nAChR functional responses in mouse MHb and IPN (Zhao-Shea et al., 2013; Shih et al., 2015; Banala et al., 2018; Arvin et al., 2019). Moreover, we recently demonstrated that continuous access nicotine SA (analogous to ContA in the present study) enhances functional nicotinic responses in MHb neuronal somata and dendrites of rats (Jin et al., 2020). The present study adds to this literature with the demonstration that somatodendritic nAChR responses in IPN neurons are enhanced following IntA nicotine SA. Coupled with the ability of IntA nicotine SA to support nicotine intake (**Fig. 7C**) and provoke relapse-like behavior (**Fig. 4**), these electrophysiology results further validate the translational relevance of the IntA nicotine SA model. Although we did not pharmacologically identify the upregulated nAChRs, some inferences can be drawn. Given that our recordings were focused on the rostral subnucleus of the IPN (i.e., IPR), where neurons express high levels of α3, β4, and α5 nAChR subunits (Shih et al., 2014; Ables et al., 2017; Arvin et al., 2019), the upregulated receptors we identified likely include those containing the β4 and α5 subunits (Hsu et al., 2013; Morton et al., 2018). However, α4β2 nAChRs cannot be excluded (Hsu et al., 2013; Shih et al., 2014). Our control comprised a group of animals that self-administered saline under IntA conditions, indicating that it is extremely unlikely for conditioned cues (rather than nicotine) to account for IPN nAChR upregulation. We were surprised to find that IPN nAChR upregulation persisted after a nicotine-free period of 7 d. Although chronic nicotine exposure can elicit very durable excitability changes (Penton et al., 2011), nAChR upregulation itself was found to reverse ∼4 days following cessation of passive nicotine exposure in mice (Marks et al., 1985). Given that ACh levels are enhanced in IPN after chronic nicotine exposure (Correa et al., 2019), coupled with the important role played by IPN neurons in nicotine withdrawal (Fowler et al., 2011; Zhao-Shea et al., 2013) and relapse (Forget et al., 2018), it will be important to investigate the role played by this persistent IPN nAChR upregulation in cue- or nicotine-elicited relapse behavior.

### Advantages of Intermittent Nicotine Self-Administration

It is well-known that drug delivery kinetics play a crucial role in drug self-administration and reinforcing efficacy (Samaha and Robinson, 2005; Allain et al., 2015). For example, nicotine is addictive when administered rapidly via a tobacco product but is therapeutic for *treating* nicotine addiction when administered slowly and continuously via a transdermal patch (Allain et al., 2015). Intermittent drug delivery promotes certain aspects of drug self-administration in animals, including nicotine (Goldberg et al., 1981; Goldberg and Henningfield, 1988). The IntA model used in this study has several advantages. First, whereas ContA permits rapid titration of plasma nicotine levels to ∼40 ng/mL in the first ∼20 min of the session, plasma nicotine levels rise more slowly during an IntA session (**Fig. 3A**). This likely enables rats to avoid an “overshoot” in brain nicotine concentration that could occur with ContA. Future studies that vary the access parameters during IntA (trial duration, ITI duration, nicotine dose, etc.) could be used to examine the relationship between estimated plasma nicotine levels and nicotine taking. Second, our data suggests that IntA is reliably able to trigger incubation of craving and therefore serves as a useful tool for investigation of nicotine relapse-like behavior (**Fig. 4** vs. **Fig. 7H,I**). Third, with the identification of a timeout as an important factor for nicotine intake (**Fig. 8**), the IntA model provides an opportunity to further investigate how cues (temporal and visual) interact with intermittent nicotine pharmacokinetics to promote nicotine intake. Finally, coupled with prior research supporting the notion that human nicotine intake can be intermittent (Henningfield et al., 1983), it is implicit that IntA nicotine SA may better model human nicotine intake for those who are not able – due to smoking restrictions imposed by cities, workplaces, or household family members – to continuously smoke/vape at any desired moment of the day.

## Limitations

Our pharmacokinetic model provides an estimate of plasma nicotine levels based on a rat’s own history of nicotine infusions, but it is limited by the fact that it does not account for individual differences in animal weight or nicotine metabolism (Kyerematen et al., 1988). Thus, although the model reports estimated plasma nicotine levels that are consistent with levels seen in prior rat studies (Craig et al., 2014) and human smokers (Benowitz et al., 1982; Benowitz et al., 1983), empirical measurements of nicotine in plasma may differ slightly from our estimates. Such empirical measurements of plasma nicotine during a labile behavioral task such as intravenous self-administration are impractical, however, as the process of blood collection could interfere with operant responding. Additionally, this study only examined male rats, precluding our ability to make specific statements or conclusions about intermittent nicotine SA in female rats.

## Conclusion

These data demonstrate that rats will readily self-administer nicotine on an intermittent access schedule and that such self-administration is sufficient to produce nAChR functional upregulation. Contingent, intermittent nicotine exposure is also sufficient to potentiate nicotine cravings. Given that many tobacco smokers and electronic cigarette users consume nicotine intermittently throughout the day, our IntA model will be useful for mechanistic investigations and for testing novel strategies to attenuate nicotine intake. Due to nicotine pharmacokinetics, intermittent interaction with combusted tobacco products may promote intake of toxic constituents compared to interaction without intermittency.

## Acknowledgments

This work was supported by National Institutes of Health (NIH) grants (DA040626 and DA035942 to R.M.D., DA048490 and DA006634 to S.R.J., and AG061787 to J.N.S.), and a pilot grant from the Wake Forest Comprehensive Cancer Center Tobacco Control Center of Excellence to K.K.S.S. and R.M.D.

## Author Contributions

M.A.T., X-T.J., K.K.S.S., S.R.J., and R.M.D. designed research, M.A.T., X-T.J., B.R.T., L.N.T., N.B.W., S.E., S.E.A., N.D., I.K., and R.M.D. performed research, M.A.T., X-T.J., N.B.W., V.J.K., J.N.S., K.K.S.S., S.R.J., and R.M.D. analyzed data, M.A.T. and R.M.D. wrote the paper.

---

**Extended Data Figure 5-1.**
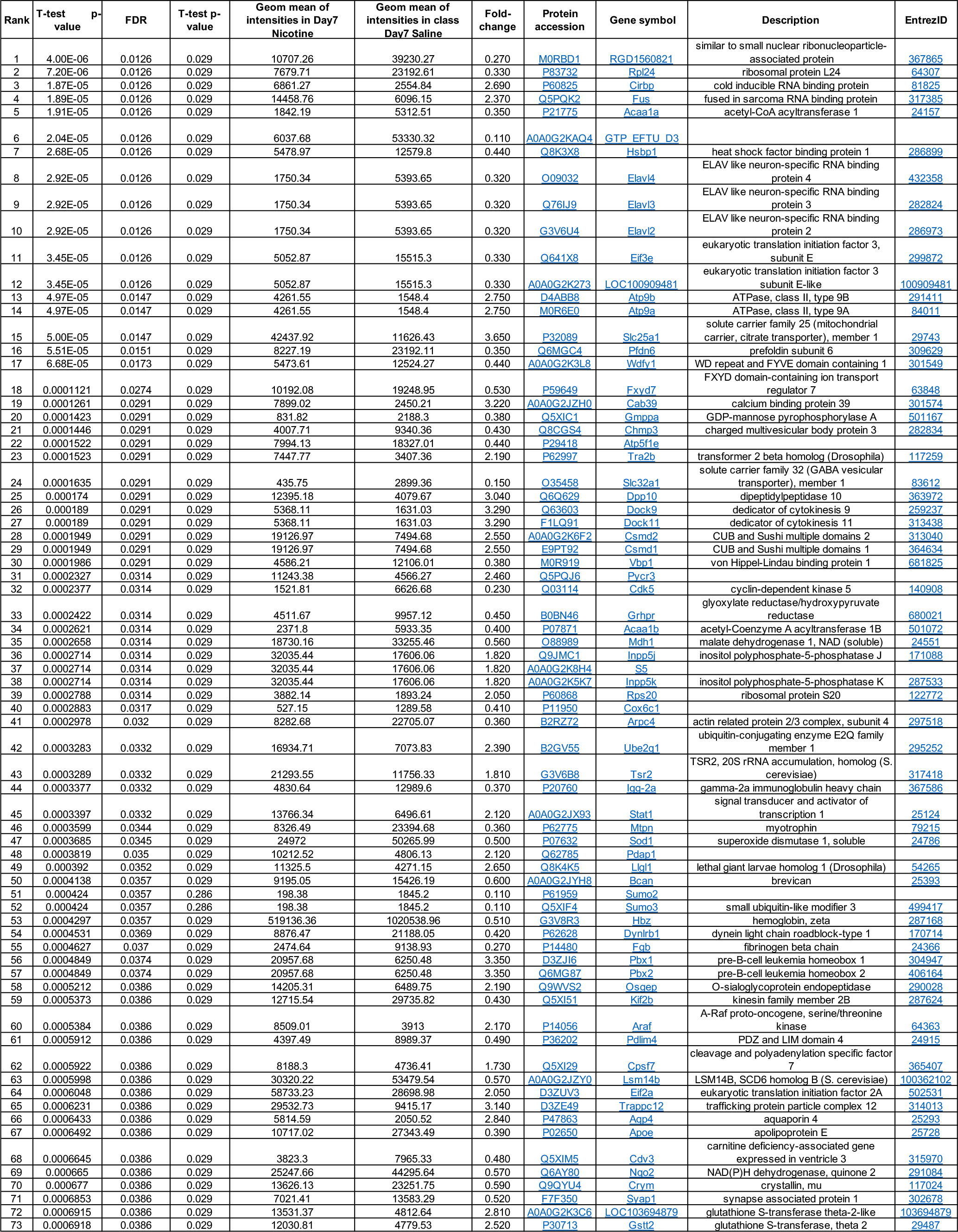
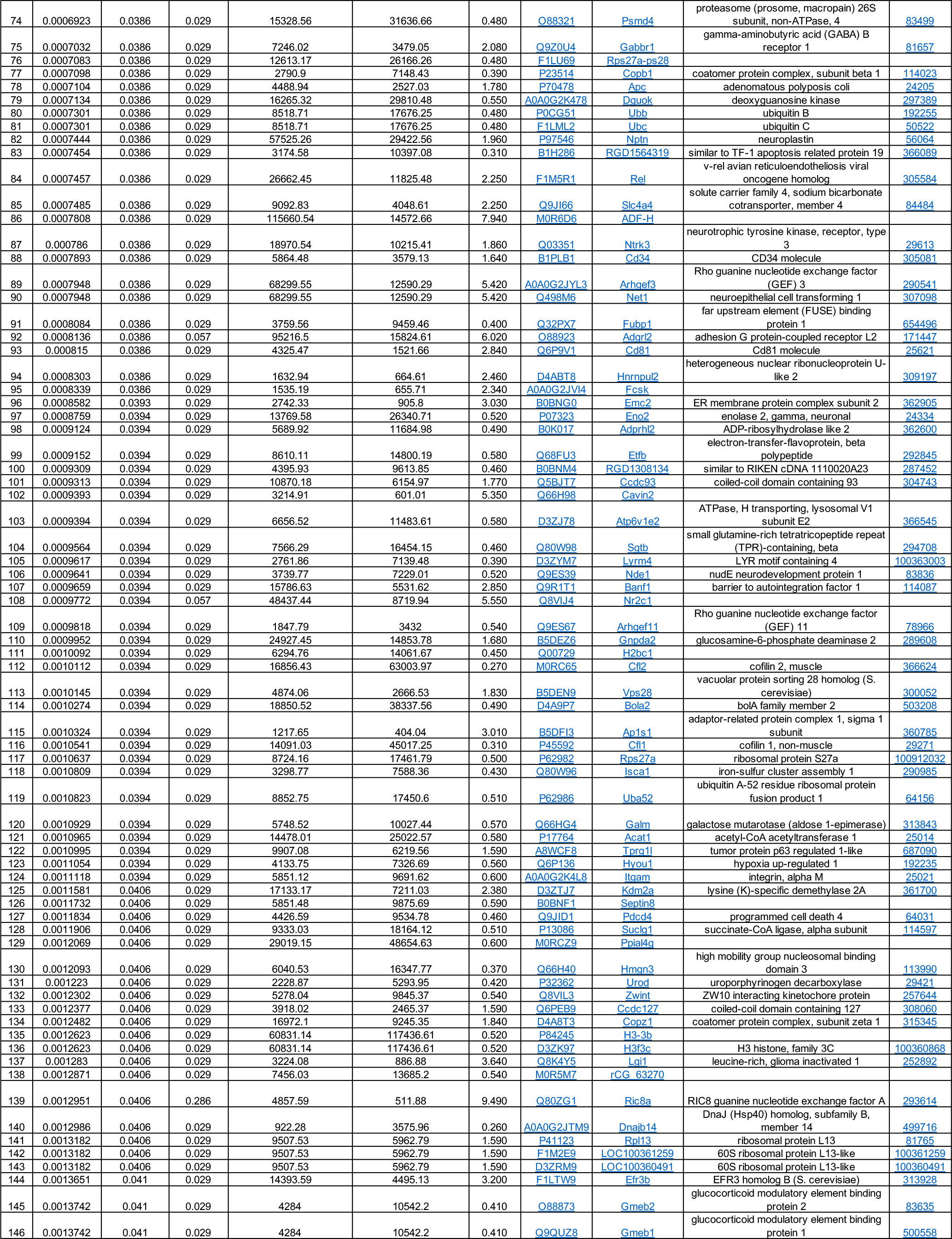
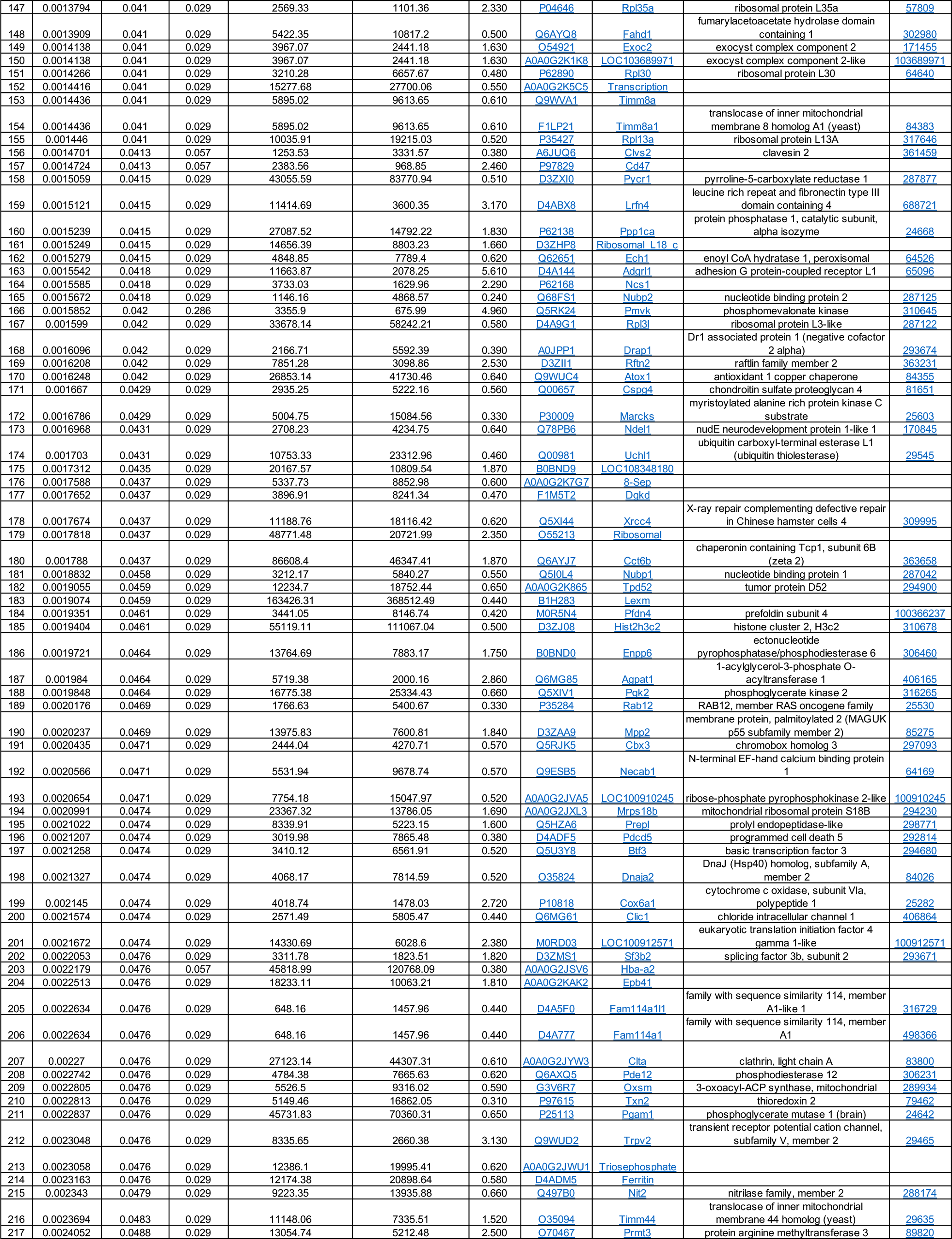
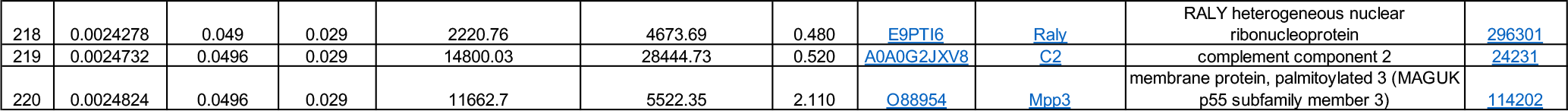
Full details for all 220 significantly altered dorsal striatum proteins are shown for nicotine SA (day 7 drug-free) vs. saline SA (day 7 drug-free).

